# Interplay between flap endonuclease 1 and DNA polymerase β in long patch base excision repair: new facets in mechanism

**DOI:** 10.64898/2026.09.22.753423

**Authors:** Yuliya S. Krasikova, Ekaterina A. Maltseva, Svetlana N. Khodyreva, Nadejda I. Rechkunova, Olga I. Lavrik

## Abstract

Mammalian base excision repair (BER) is the major repair pathway for correcting small DNA base lesions. BER operates via two sub-pathways: short patch (or single nucleotide) and long patch BER (SP-BER and LP-BER, respectively). In LP-BER, DNA polymerase beta (Polβ) performs strand-displacement synthesis generating a 5’-flap that is removed by flap endonuclease 1 (FEN1). To determine the precise DNA-intermediate that “switches on” FEN1 activity, we analyzed the FEN1 cleavage position in DNA-intermediates generated by Polβ. For simultaneous detection of Polβ-catalyzed primer elongation and FEN1’s cleavage products, we utilized DNA structures with fluorescent labels on the upstream and downstream primers. Our data indicate that FEN1 operates efficiently on DNAs containing one or more equilibrium nucleotides. FEN1 binds to the 3’-terminal equilibrium nucleotide of the primer and probes it for a 3’-OH group. The cleavage point is located directly opposite this 3’-terminal equilibrium nucleotide of the primer. The subsequent elimination of the equilibrium area leads to restoration of the ligatable structure, thereby limiting further strand-displacement synthesis. Since the product sets generated by the two enzymes are mirror-symmetrical, we propose that pausing of Polβ permits FEN1 to join the Polβ-DNA complex. Our study demonstrates that the FEN1-Polβ interplay occurs via the generation and elimination of equilibrium nucleotide areas rather than through a “gap-translation” mechanism. The length of the equilibrium area depends on dNTP concentration, suggesting that the size of the LP-BER repair patches may differ between S-phase and the remainder of the cell cycle.

**Highlights:**

FEN1 operates on DNAs with equilibrium nucleotides inserted by Polβ.
FEN1 binds the 3’-terminal equilibrium nucleotide and probes it for a 3’-OH.
The cleavage point is located directly opposite the 3’-end of the equilibrium area.
FEN1-Polβ interplay occurs via the generation and elimination of the equilibrium area.
DNA is channeled from Polβ to FEN1 within the ternary complex.

**Graphical abstract:** 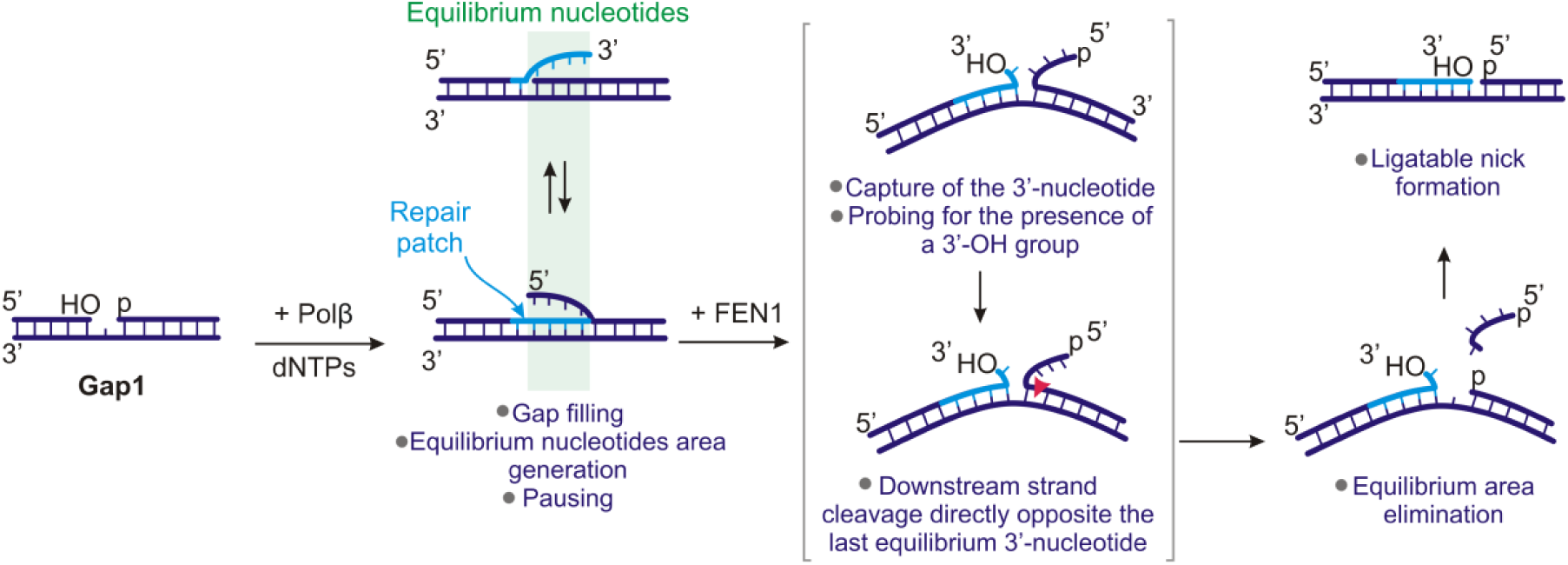

## 1. Introduction

Base excision repair (BER) is the predominant repair mechanism responsible for correcting non-helix-distorting base lesions produced by oxidation, alkylation, or deamination [1], [2], [3]. This repair process is initiated by DNA glycosylases, which recognize and remove the damaged base, generating an apurinic/apyrimidinic (AP) site. In eukaryotic cells, the major enzyme that catalyzes cleavage of AP sites is AP endonuclease 1 [4], [5]. This results in the formation of a 3’-OH group and a 5’-deoxyribose phosphate (dRP)-residue at the edges of the one nucleotide gap. DNA polymerase β (Polβ) then removes the dRP-residue via its 5’-lyase activity and fills the gap [6], [7], [8], generating a nick that is sealed by the DNA ligase III-XRCC1 complex [9], [10], [11]. This BER sub-pathway is designated a short patch (SP-BER) or single-nucleotide base excision repair (SN-BER), owing to the one nucleotide repair patch.

Alternatively, if nick ligation is restrained for some reason, such as a low ATP concentration or end modification, Polβ continues synthesis with the downstream strand-displacement, switching the repair process to the long-patch sub-pathway (LP-BER) [12]. Commonly, LP-BER operates when the 5’-dRP-residue cannot be removed by the Polβ lyase activity for the reason of oxidation or reduction. In this case, Polβ instead performs strand-displacement synthesis, thereby producing a 5’-flap [16]. LP-BER is also proposed to be the dominant mechanism of the post-replicative BER initiated by bi-functional DNA glycosylases [13], [14], [15]. Flap endonuclease 1 (FEN1) then cleaves the DNA strand containing the flap [17], [18], [19], and the resulting nick is subsequently sealed by DNA ligase I (Lig1) [20], [21]. Thus, in the LP-BER process, FEN1’s main partners are the upstream operating Polβ and the downstream operating Lig1.

The interaction between FEN1 and Polβ has been the subject of extensive research for decades. Previous studies have demonstrated that Polβ stimulates FEN1 excision activity [22] and that these enzymes interact physically [23], [24]. Formation of the ternary Polβ-FEN1-DNA complex was not detected, but FEN1 co-immunoprecipitated with photoreactive-DNA-crosslinked Polβ [25], and Polβ activated cleavage by FEN1 trapped on the flap-DNA [24]. Nevertheless, DNA handover within the ternary complex has been proposed as the “passing the baton” LP-BER mechanism. Alternatively, Polβ may create a DNA intermediate that FEN1 can recognize and process independently (the “hit-and-run” LP-BER mechanism [26], [27]). Moreover, according to a previous finding, Polβ engages in cooperation with FEN1 through the “gap-translation” process: Polβ fills an initial 1nt gap, displacing an altered dRP-group into the 5’-flap, FEN1 then cleaves it at the -1 nt position in the duplex part, generating 1nt gap, Polβ then fills that gap, converting it to a nick and generating a substrate for FEN1, whose cleavage results in a new 1nt gap – a substrate for Polβ, *et cetera* [16], [26], [27]. Therefore, alternating Polβ gap filling and FEN1 gap generation results in “gap translation”. *In vivo* experiments have revealed that at least half of BER events are processed through the LP-BER sub-pathway [28]. In particular, the mean length of the repair patches was 2–6 nucleotides for 8-oxo-7,8-dihydroguanine and 6–12 nucleotides for 3-hydroxy-2-(hydroxymethyl)-tetrahydrofuran (a synthetic analog of the AP-site). Subsequently, using HEK293 cell extracts, the authors of [29] showed that these cells had Polβ-dependent BER (owing to its essential resistance to aphidicolin) with mostly 2nt repair patches.

FEN1 is a highly conservative structure-specific nuclease that processes intermediates of DNA replication and repair when process-associated polymerases perform strand-displacement synthesis, producing single-stranded (ss) 5’-flaps at junctions with double-stranded (ds) DNA. Besides LP-BER, FEN1 participates in Okazaki fragment maturation, telomere maintenance, and processing of stalled replication forks [30], [31], [32].

It was found earlier that FEN1 preferentially binds double-flap substrates with a 1nt 3’-flap and a 5’-flap of any length, including zero [33], [34], [35]. These structures mimic equilibrium replication intermediates [36], but it is not clear how FEN1 can access the 3’-nucleotide of a primer. In fact, the 3’-nucleotide is tightly bound by the polymerase, and there is no known mechanism of polymerase backtracking to make space for FEN1’s interaction; hence, it must be either a mechanism of direct handover or one of polymerase dissociation from the DNA. Later, it was discovered that during replication, Polδ-FEN1 switching occurs via the PCNA toolbelt [37]. What mechanism operates during PCNA-independent LP-BER [16], [38] is still unclear. With that, several questions arise: which LP-BER DNA-intermediate “switches on” FEN1 cleavage activity, and whether repair patches of about 12 nucleotides in length are produced one by one through a number of filling-cleavage steps, or by a single cleavage event on an extended 5’-flap?

Here, to address these questions, we examined the position of the FEN1 cleavage point on LP-BER DNA-intermediates that were preformed or generated by Polβ. For this purpose, we designed DNA structures modeling BER intermediates and performed *in vitro* biochemical experiments. Electrophoretic separation of the reaction products allowed distinguishing single-nucleotide resolution, revealing the precise FEN1 cleavage point. Each DNA contained fluorescent labels on the upstream and downstream primers, enabling detection of Polβ primer extension products and FEN1 cleavage products within a single reaction mixture. We found that FEN1 operates on DNA intermediates containing at least one equilibrium nucleotide generated by Polβ. Equilibrium nucleotide(s) are the nucleotide(s) inserted by Polβ through strand-displacement synthesis which are complementary to the same nucleotide(s) in the template as the displaced nucleotides of the downstream primer. FEN1 interacts with the 3’-terminal Polβ-inserted nucleotide and probes it for the presence of a 3’-OH group. The cleavage point of the 5’-flap strand is located directly opposite this terminal equilibrium 3’-nucleotide of the primer. The subsequent elimination of the equilibrium area leads to restoration of the ligatable structure, thereby limiting further strand-displacement synthesis. Since the product sets generated by the two enzymes are mirror-symmetrical, we propose that Polβ pausing allows FEN1 to cleave the flap. Exploring the possibilities for Polβ-FEN1-DNA complex formation, we discovered that the ternary complex forms with both substrate and product DNAs. The length of the equilibrium area depends on dNTP concentration, suggesting that the size of the LP-BER repair patches may differ between S-phase and the rest of the cell cycle.

## 2. Materials and Methods

### 2.1 Reagents and Oligonucleotides

BSA was from Sigma (USA); reagents for electrophoresis and buffer components were acquired from either Sigma (USA) or Russian vendors (extra-pure grade). Oligonucleotides bearing a fluorescent label FAM or TAMRA were custom-synthesized by the center for high-precision DNA/RNA synthesis of ICBFM SB RAS or by Lumiprobe (Russia). Oligonucleotide sequences and the resulting DNA structures are listed in the Supplementary Materials (**Tables S1** and **S2**).

### 2.2 Preparation of DNA Structures

DNA structures were annealed by mixing (template): (FAM-labeled oligonucleotide for 5’-flap strand): (non-labeled or TAMRA-labeled primer s) in 1:1:1 molar ratio in a <u>PB</u> <u>buffer</u> solution containing 50 mM Tris-HCl (pH 7.5), 1 mM DTT, and 100 mM KCl [39]. The reaction mixtures were incubated for 3 min at 95 and then cooled slowly to room temperature. The degree of hybridization was monitored by electrophoresis in a 10% polyacrylamide gel (acrylamide/bis-acrylamide = 40:1) in TBE buffer (89 mM Tris, 89 mM H_3_BO_3_, 2 mM EDTA, pH 8.3). All DNA structures were obtained in high yield and were stable under these conditions.

### 2.3 Protein Purification

The pET28-FSH plasmid containing human FEN1 cDNA was kindly provided by Dr. R. Prasad (National Institute of Environmental Health Sciences, USA). Recombinant human FEN1 was produced in *Escherichia coli* and purified essentially as described in [40], [41]. Briefly, *E. coli* BL21(DE3) was transformed with the pET28-FSH plasmid, grown at 30°C, and FEN1 synthesis was induced with IPTG at a final concentration of 0.5 mM for 4 hours at 25°C. Cells were collected by centrifugation and lysed. DNA was fragmented by sonication, followed by removal of cellular debris. Recombinant human FEN1 (hFEN1), containing a His-tag at the C-terminus, was purified using several chromatographic steps: chromatography on Ni-NTA agarose, SP-Sepharose, and heparin-Sepharose. The resulting hFEN1 sample had a purity of >95%, as determined by sodium dodecyl sulfate-polyacrylamide gel electrophoresis [42] and Coomassie staining. The purified enzyme was stored in buffer (40 mM sodium phosphate buffer, pH 7.8, 50% glycerol, 0.1 M NaCl, 0.1% NP-40, 1 mM EDTA, and 5 mM DTT) at -20°C.

Plasmid pRSETB-pol, containing the full-length cDNA of the rat *POLB* gene, was a generous gift from Samuel H. Wilson (National Institute of Environmental Health Sciences, Galveston, USA). Recombinant Polβ was overexpressed in the Rosetta 2(DE3)pLysS strain of *E. coli* (Novagen, Germany) and purified as previously described [43].

### 2.4 Enzymatic Assay

FEN1 nuclease activity is highly sensitive to salt concentration, so the KCl concentration in all reaction mixtures was adjusted to 100 mM (except in the experiments examining the salt dependence of FEN1 exonuclease activity, where the total salt concentration was adjusted to 15, 50, 100, or 150 mM). For this purpose, we mixed reactions with 50 mM KCl and then added protein dilution buffer (buffer PB with 0.6 mg/mL BSA) in the amount required to obtain a final concentration of 100 mM KCl, taking into account the salt from the protein aliquots. Thus, reaction mixtures containing buffer PB, at least 0.6 mg/mL BSA (BSA amount increased due to the added volume of protein dilution buffer), 5 mM MgCl_2_, 0.5 mM MnCl_2_, 20 nM DNA, 20 nM FEN1 (unless otherwise stated), 20 nM Polβ (unless otherwise stated), as well as 1 or 50 µM dNTPs (dTTP/ddTTP, dATP/ddATP, dCTP, dGTP), were assembled on ice.

Reactions were initiated by transferring the reaction mixtures to 37°C and incubating them for the periods indicated in the figure legends. For kinetic experiments, 8 or 9 µL of the reaction mixture was taken at various time intervals. For the protein titration experiments, the reaction mixtures were incubated for 30 minutes. The reactions were terminated by adding 8 µL of hot formamide (95°C) and heating the mixtures to 95°C. These samples were then loaded onto a heated (approximately 45 °C) denaturing 10% or 20% polyacrylamide gel (acrylamide/bis-acrylamide = 19:1, 8 M urea, and 10% or 20% formamide) and separated by electrophoresis in TBE buffer. The reaction products were detected in FAM or TAMRA mode using a Typhoon FLA 9500 scanner (GE Healthcare). Scanned images were visualized and analyzed using Quantity One software. The experiment with each DNA structure was performed in at least triplicate.

### 2.5 EMSA

Reaction mixtures (10 µL) consisting of PB buffer, 0.6 mg/mL BSA, 20 nM DNA, 5 mM MgCl2 or 5 mM EDTA, 10 µM dNTPs (where indicated), 200 or 400 nM FEN1 (unless otherwise stated), and 100 or 200 nM Polβ (unless otherwise stated) were assembled on ice. Reactions were initiated by transferring the reaction mixtures to 37 °C and incubating them for 10 minutes. The protein-bound complexes were then immediately placed on ice and 2 µL of loading buffer was added (9% Ficoll, 0.015% bromophenol blue in PB buffer with 0.6 mg/mL BSA). Next, the samples were loaded onto pre-run (about 2h) 7% native polyacrylamide gel (acrylamide/bis-acrylamide = 40:1) and separated by electrophoresis in 1×TBE at 4 °C with a voltage decrease of 8-10 V/cm. The gels were than scanned between glass plates in FAM-mode on a Typhoon FLA 9500 (GE Healthcare) and visualized using Quantity One software.

## 3. Results

### 3.1 The position of the FEN1 cleavage point is independent of the 5’-flap length

The ability of FEN1 to catalyze the removal of 5’-flaps from hybrid single-stranded/double-stranded DNA structures has been the subject of extensive research employing a variety of techniques over the past two decades. According to structural data, FEN1 hydrolyzes a phosphodiester bond in the 5’-flap strand precisely one nucleotide into the duplex [17], [18], [19]. Indeed, given the order of events in the BER and Okazaki maturation processes, FEN1 hands off the DNA to DNA ligase 1, so correct nick formation is indispensable for ligation. However, numerous biochemical experiments with FEN1 have yielded more than one cleavage product (e.g. [44], [45]). Discrepancies between the hypothesis and the experimental data could potentially be attributed to the influence of salt concentration (typically 50 mM or less) and flap length. To clarify this, we performed experiments with DNAs containing flaps of different lengths (1, 2, 3, or 5 nucleotides) coupled with a nick under conditions close to those found in body cells (see **Fig. 1 and Table S2**).

**Fig. 1.**
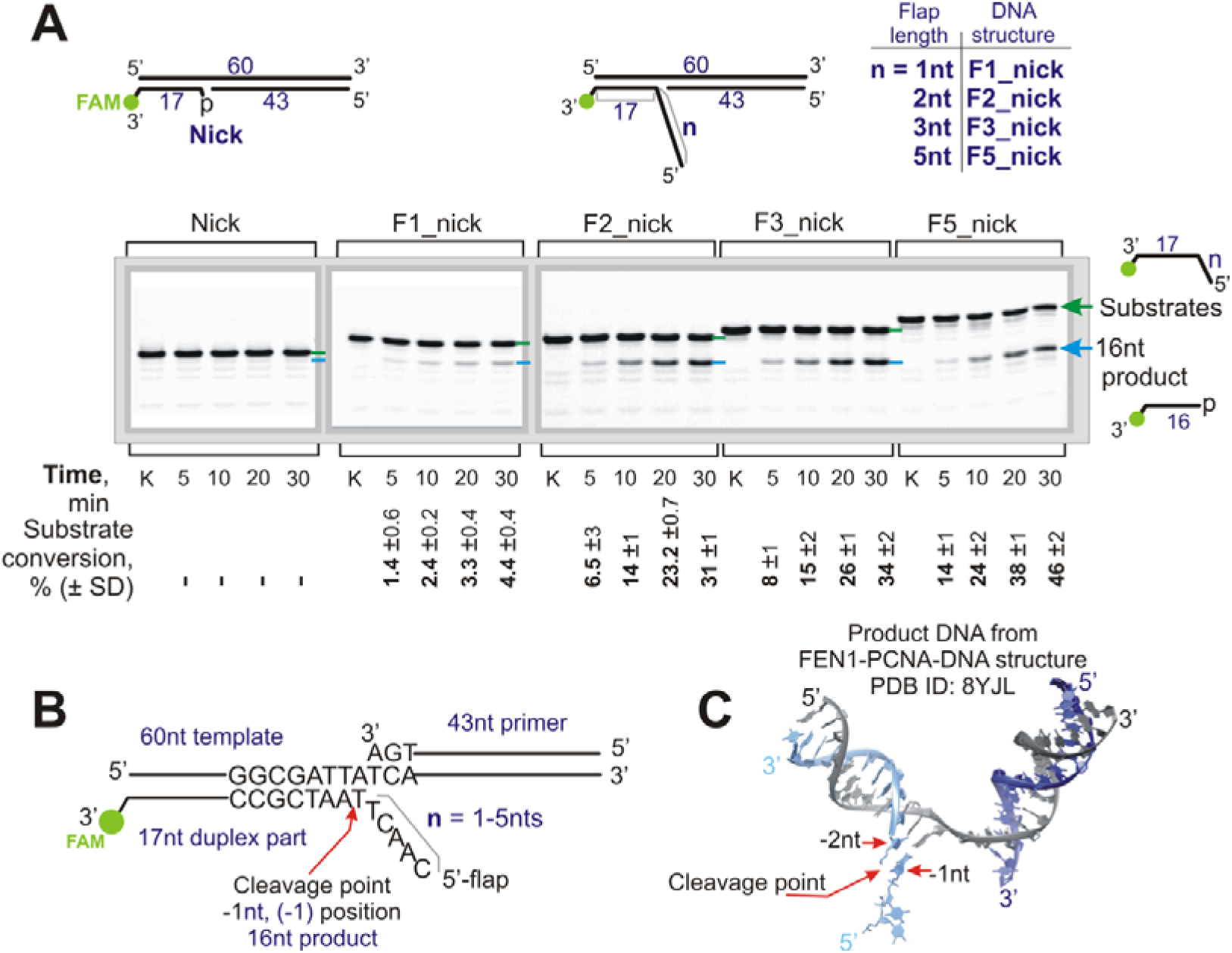
FEN1 cleaves flap-containing DNAs at a defined position regardless of flap length. FEN1 cleaves DNA-structures containing a flap 1, 2, 3, or 5nt coupled with a nick at the same position. (**A**) The cleavage products were 16nt in length, regardless of the flap length. These products were produced by cleavage in the duplex part and corresponded to the (-1) position from the branch point. Nicked DNA was not cleaved. A schematic representation of the substrates is depicted at the top; the lime green circle indicates FAM-label position; green arrows/lines indicate the position of the substrate DNA; blue arrows/lines indicate the product DNA. Lane “k” is a DNA control. The reaction mixtures contained 20 nM DNA and 20 nM FEN1. The percentage of substrate conversion indicates the conversion of substrate to product. (**B**) Schematic representation of the structure of the DNAs used with cleavage point indicated. FEN1 cleaves the flap-strand in the duplex part one nucleotide downstream of the first paired nucleotide at the (-1) position from the branch point. The flaps in the substrate DNAs formed because they were not complementary to the template sequence. (**C**) The product DNA image was obtained from the cryo-EM structure of the FEN1-PCNA-DNA complex, PDB ID: 8YJL. This DNA was endogenous and was co-purified from HEK293 cells transfected with a plasmid encoding affinity-tagged PCNA [19]. It is clearly seen that there is a break in the density between the first (-1nt) and second (-2nt) nucleotides.

Our experiments demonstrate that FEN1 cleaves all DNAs at the same position regardless of the 5’-flap length (**Fig. 1A**). The cleavage products were 16nt long and corresponded to the (-1) position from the branch point in the duplex part. After 5’-flap elimination, the product DNA contains a 1nt gap bearing a phosphate at the 5’-margin. Mapping the cleavage point onto the DNA sequence revealed that FEN1 cleaves the flap strand at the phosphodiester bond located after the first paired nucleotide within the 17nt duplex region (**Fig. 1B**). FEN1 demonstrates the same cleavage position with DNA containing synthetic analogues of 5’-dRP (tetrahydrofuran phosphate, THFp) as well **(Fig. S1**). This is fully consistent with cryo-EM structural data indicating that cleavage occurs at the phosphate between the -1 and -2 nucleotides adjacent to the branch point (**Fig. 1C**) [19].

FEN1 does not process nicked DNA (**Fig. 2A**). This result could be expected since nick DNA is ligatable and does not require FEN1 processing. However, this result contrasts with literature data on FEN1 exonuclease activity detected on nick DNA [46]. Furthermore, it has previously been reported that FEN1 can cleave the flap strand in a step-by-step manner [16], [47]. We propose that the exonuclease activity may be due to a low concentration of monovalent metal ions and a high concentration of FEN1. In other words, low-salt conditions decrease duplex stability and facilitate duplex accommodation to the catalytically active FEN1-DNA complex. Subsequently, an increase in FEN1 activity could result in several cleavage points. To test this proposition, we examined the relationship between the number of FEN1 cleavage products, enzyme concentration, and ionic strength (**Fig. S2**).

**Fig. 2.**
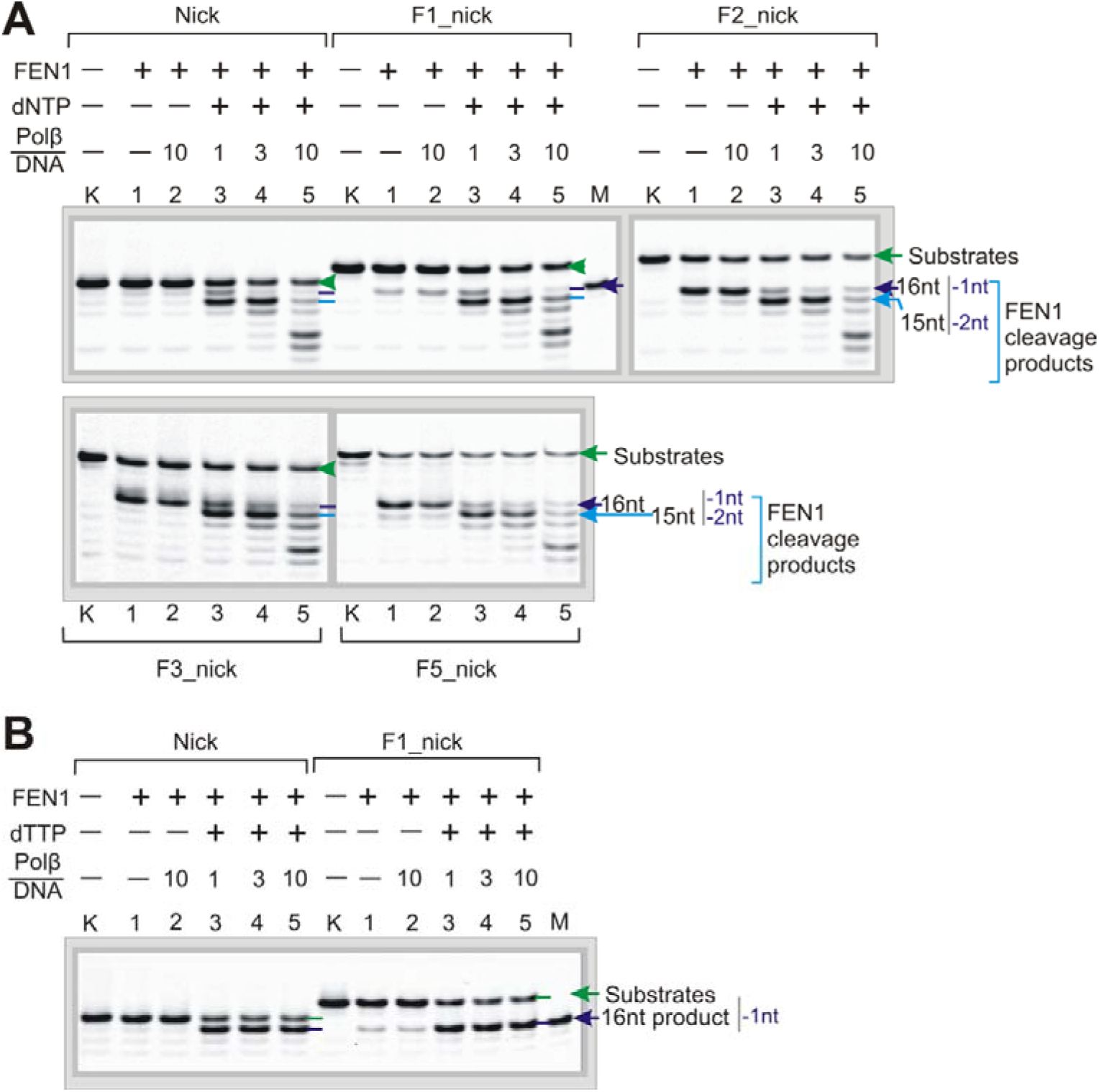
The FEN1 cleavage point shifts to the (-2) position in the presence of Polβ and all dNTPs (**A**) and remains at the (-1) position in the presence of dTTP (**B**). The DNA substrate names are indicated at the top; green arrows/lines indicate the position of the substrate DNA; dark and light blue arrows/lines indicate the cleavage products of 16nt and 15nt, respectively. Lane “k” is the DNA control; lane “M” is the length marker. Quantitative data for the (A) experiments are presented in Fig. S3. Reaction mixtures contained 20 nM DNA, 20 nM FEN1, 1 µM dNTPs or dTTP (unless otherwise stated), and Polβ at the indicated enzyme/DNA ratios. Reaction products were detected in FAM-mode.

When the reactions were carried out at physiological salt concentrations (100-150 mМ) and with an equimolar concentration of FEN1 and DNA, cleavage of nicked DNA was not observed. Products of exonuclease cleavage appeared at high excess of FEN1 over DNA, and were clearly detectable at all enzyme concentration used when salt concentrations did not exceed 50 mM. Notably, nicked DNA containing a non-complementary 3’-flap was effectively cleaved at the (-1) position, even at a salt concentration of 150 mM. Decreasing the salt concentration and increasing the FEN1:DNA ratio subsequently led to the appearance of several additional cleavage points.

From the standpoint that the predominant LP-BER product is 2nt [29], we expected that FEN1 would efficiently cleave flaps consisting of 1nt or a dRP-group (natural or oxidized/reduced). Unexpectedly, our experiments revealed that FEN1 cleaves such flaps with very low efficiency (Fig. 1A and Fig. S1). Moreover, cleavage efficiency increases with increasing flap-length. We found literature data with the same result [48]. Therefore, we further decided to test the possible modulating influence of Polβ on the FEN1’s ability to cleave 1nt-flaps. Especially since Polβ is one of FEN1’s main interacting partners in the BER process.

### 3.2 Polβ provides conditions for FEN1-dependent cleavage of nicked DNA and facilitates a shift of the flap-strand cleavage point to the (-2) position relative to the initial branch point

It has been proposed that subsequent synthesis by Polβ to add a nucleotide in the space occupied by an oxidized/reduced dRP-group creates an intermediate that resists further synthesis and thus Polβ promotes the removal of a forward barrier by FEN1 [26]. Thereby, we hypothesized that Polβ could stimulate not only overall FEN1 cleavage efficiency but also facilitate flap-strand cleavage at a defined position. To test this hypothesis, we investigated the relationship between cleavage position and Polβ activity and flap length (**Fig. 2A**). All reactions were carried out in the presence of 1 µM dNTPs, which corresponds to the concentration of deoxy nucleotides throughout the cell cycle, excluding S-phase [49]. As expected, Polβ stimulates the cleavage of all flap-containing DNAs by FEN1. Specifically, Polβ increases the efficiency with which FEN1 cleaves nick DNA to a level comparable with that of other DNAs, and significantly increases the yield of the F1-nick product (**Fig. S3**). These results are consistent with the cell extracts data that FEN1 should cleave 1nt-long flaps with high efficiency [29]. The cleavage efficiency of 2-, 3-, and 5nt-flap DNAs increased moderately.

Unexpectedly, the cleavage point shifted towards the (-2) position at concentrations of Polβ close to equimolar, and towards the (-5) position at a 10-fold excess of Polβ. These results are consistent with the lengths of repair patches 2-6nt observed during the correction of 8oxoG by the LP-BER process in the HCT116 mammalian cell line [28]. Additionally, it coincidentally matches the gap size (2-6nt) that Polβ is capable of filling through processive DNA synthesis [50]. In the case of nicked DNA, the cleavage at the (-2) position could correspond to a single FEN1 cleavage event after 2nt primer extension by Polβ or, alternatively, to two cycles of filling-cleavage. Thereafter, we asked whether these flap-cleavage products at the (-2) position were produced through a single cleavage-filling event.

To determine the number of cleavage events corresponding to a single-nucleotide primer extension, experiments were performed in the presence of dTTP only (**Fig. 2B**). The results indicated that FEN1 initiates cleavage of the nick DNA and that cleavage of a 1nt-flap DNA is significantly improved (compare lanes 1-2 with 3-5 for the corresponding DNAs). The resulting product corresponds to one cleavage event at the (-1) position. This result can be explained by two scenarios. First, Polβ stimulates FEN1 cleavage through protein-protein interactions, creating an ideal substrate for gap filling catalyzed by Polβ [26]. However, the question remains why Polβ does not induce further FEN1 cleavage events and why the “gap/nick translation process” is not observed? The second scenario implies that Polβ extends the primer by1nt thereby producing a 1nt equilibrium 3’-flap that is a better substrate for FEN1 cleavage [24]. However, this scenario does not explain why the cleavage point shifts directly to the (-2) position in the presence of all dNTPs.

### 3.3 The FEN1 catalytic reaction is triggered by the incorporation of a correct nucleotide by Polβ

In contrast to a previous observation [24], our results demonstrate no stimulatory effect in the absence of dNTPs (lanes 2 in **Fig. 2**). Thus, it is possible that protein-protein interactions occur when Polβ transitions to the “closed” conformation in the presence of dNTP [7].

To address this question, we introduced a fluorescent TAMRA label at the 5’-end of the primer, enabling us to assess both Polβ-catalyzed primer extension and FEN1-catalysed cleavage simultaneously (**Fig. 3**). Under these reaction conditions, Polβ does not catalyze the insertion of a non-complementary nucleotide into nicked DNA (lanes 7-18 in **Fig. 3A**) and FEN1-catalyzed cleavage is not observed. In the presence of the correct dTTP, Polβ extends the primer by one nucleotide (+1) (TAMRA detection), while FEN1 cleaves the downstream primer at the (-1) position (FAM detection). The cleavage stimulation effect on 1nt-flap DNA is only observed in the presence of the correct nucleotide (**Fig. S4**).

**Fig. 3.**
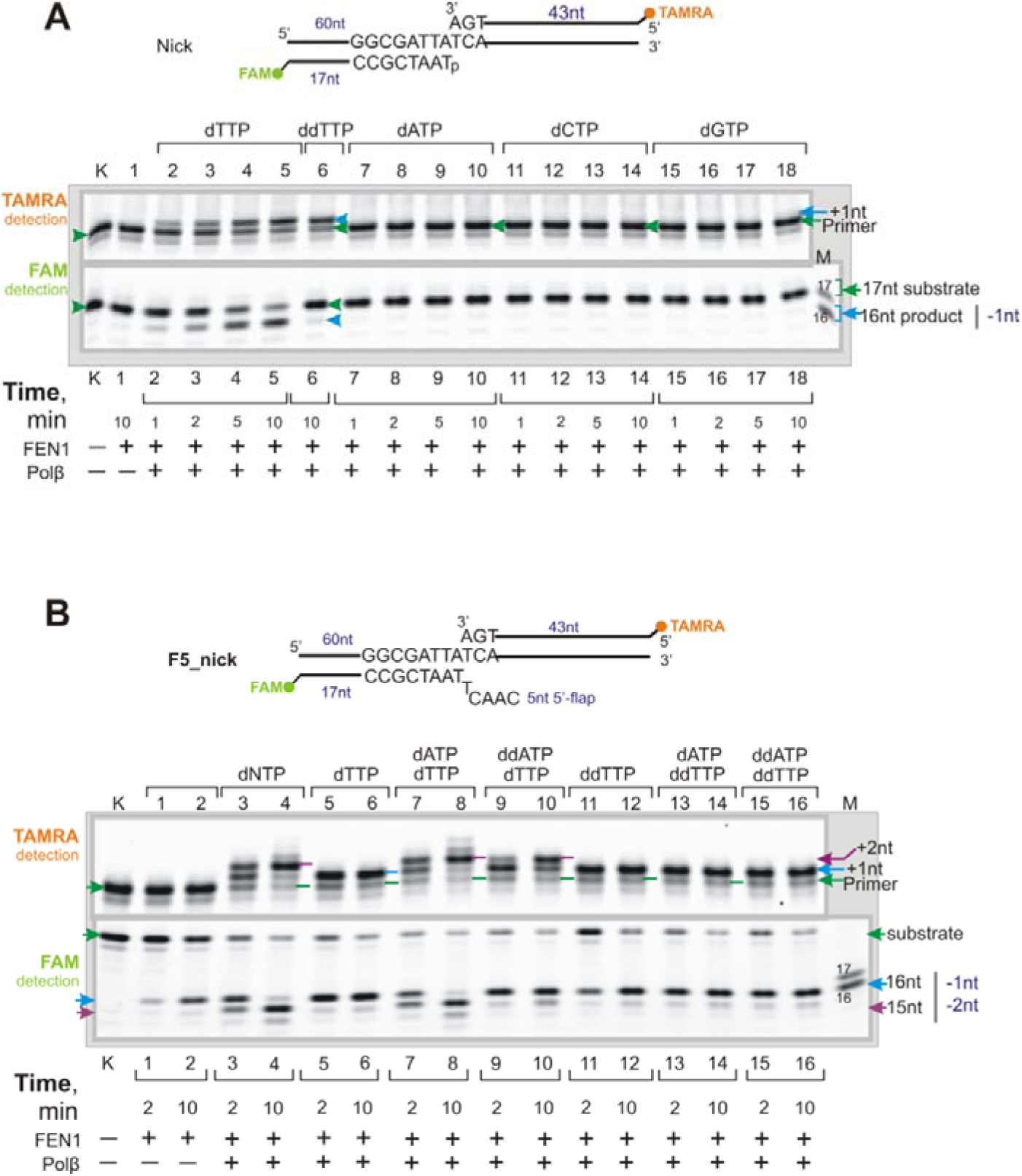
FEN1-catalyzed DNA cleavage was triggered by Polβ-mediated primer extension in nicked DNA (**A**) and 5nt-flap DNA (**B**). Products of DNA cleavage catalyzed by FEN1 were detected by a FAM label at the 3’-end of the downstream primer, and primer extension catalyzed by Polβ was detected by a TAMRA label at the 5’-end of the primer. Substrate names and schematic representations are shown above the gels; lime green and orange circles denote FAM and TAMRA labels, respectively; green arrows/lines indicate the position of substrate DNA (primer or flap-strand); blue arrows/lines indicate elongation or cleavage products. Lane “k” is the DNA control; lane “M” is the length marker. Reaction mixtures contained 20 nM DNA, 20 nM FEN1, 20 nM Polβ, and 1 µM dNTPs or ddNTPs (all four, or separately, or in the indicated combination).

Experiments with 5nt-flap DNA revealed that FEN1 cleavage efficiency increased in the presence of the correct dTTP compared to conditions without Polβ (compare lanes 1-2 with 5-6 in **Fig. 3B**). The cleavage effectiveness was similar in the presence of all dNTPs and in the presence of dTTP alone (lanes 3-6 in **Fig. 3B**). However, after 10 minutes, the main cleavage product for all dNTPs corresponded to the (-2) position (lane 4, FAM detection), which completely mirrored the 2nt primer elongation by Polβ. After 2 min, the cleavage products corresponded to the (-1) and (-2) positions (lane 3, FAM detection). During this time, Polβ inserts 1 or 2 nucleotides (lane 3, TAMRA detection), and the yield of the +2nt extension product matches that of the (-2) position cleavage product (lane 3, FAM detection). The resulting cleavage efficiency at the (-1) position was due to a combination of two cleavage situations: cleavage before primer extension and cleavage after 1nt primer extension. Due to the absence of stimulation in the presence of non-complementary dNTPs, we assume that the stimulation effect is not solely mediated by protein-protein interactions but rather by the formation of a DNA intermediate.

### 3.4 FEN1 cleavage activity is more dependent on the presence of a 3’-OH group at the 3’-end of the primer than on the complementarity of the 3’-nucleotide

The stimulation effect was significantly reduced when ddTTP was used as the substrate compared to dTTP (compare lanes 5-6 with 11-12 in **Fig. 3B**). The same result was obtained with 1nt-flap DNA (**Fig. S4**); however, nicked DNA was not cleaved at all (lane 6 in **Fig. 3A**). The forward nucleotide sequence in the template is ATTA and, unexpectedly, the cleavage point did not shift to the (-2) position in the presence of (dTTP+ddATP). Only a minor corresponding product became detectable (compare lanes 9-10 with 7-8 in **Fig. 3B**, as well as lanes 11-13 in **Fig. S5A** and lanes 12-15 in **Fig. S5B**). These results clearly demonstrate that both nucleotide insertion and the 3′-OH group at the 3′-end of the primer are indispensable for FEN1 cleavage stimulation.

In the crystal structure of the FEN1-DNA complex, the 3’-OH group forms hydrogen bonds with the backbone carbonyl of Lys314 and the hydroxyl group of Thr61 (**Fig. S6A**) [17]. Combining these structural data with our biochemical results, we propose that the absence of the 3’-OH group leads to incorrect positioning of the 3’-nucleotide inside the binding pocket (see **Fig. S6B**), thereby dramatically reducing the stimulatory effect.

It is noteworthy that Polβ-mediated nucleotide(s) insertion into nick-DNA results in the formation of an equilibrium region consisting of nucleotides from both upstream and downstream of the primers that are complementary to the same portion of the template. Consequently, the FEN1 natural substrate should contain an equilibrium nucleotide. This assumption was previously proposed from the standpoint of the observed stimulation effect in the presence of a 3’-flap [24], [36], [51].

Following this idea, we examined FEN1 cleavage activity using nick-DNA containing non-complementary 3’-flap and a fully complementary 3’-flap forming a 1nt equilibrium area (**Fig. 4**). FEN1 processed both substrates, cleaving the non-complementary 3’-flap about 10% more efficiently than the equilibrium one (compare lanes 3 and 5 at 5 min for nick_1eq and nick_f1). Structural data show that the 3’-nucleotide is displaced from the duplex by the hydrophobic wedge (**Fig. S6C**) and, accordingly, if this nucleotide is already unpaired, it is more readily positioned into the 3’-flap binding pocket (**Fig. S6B**) [17], [19].

**Fig. 4.**
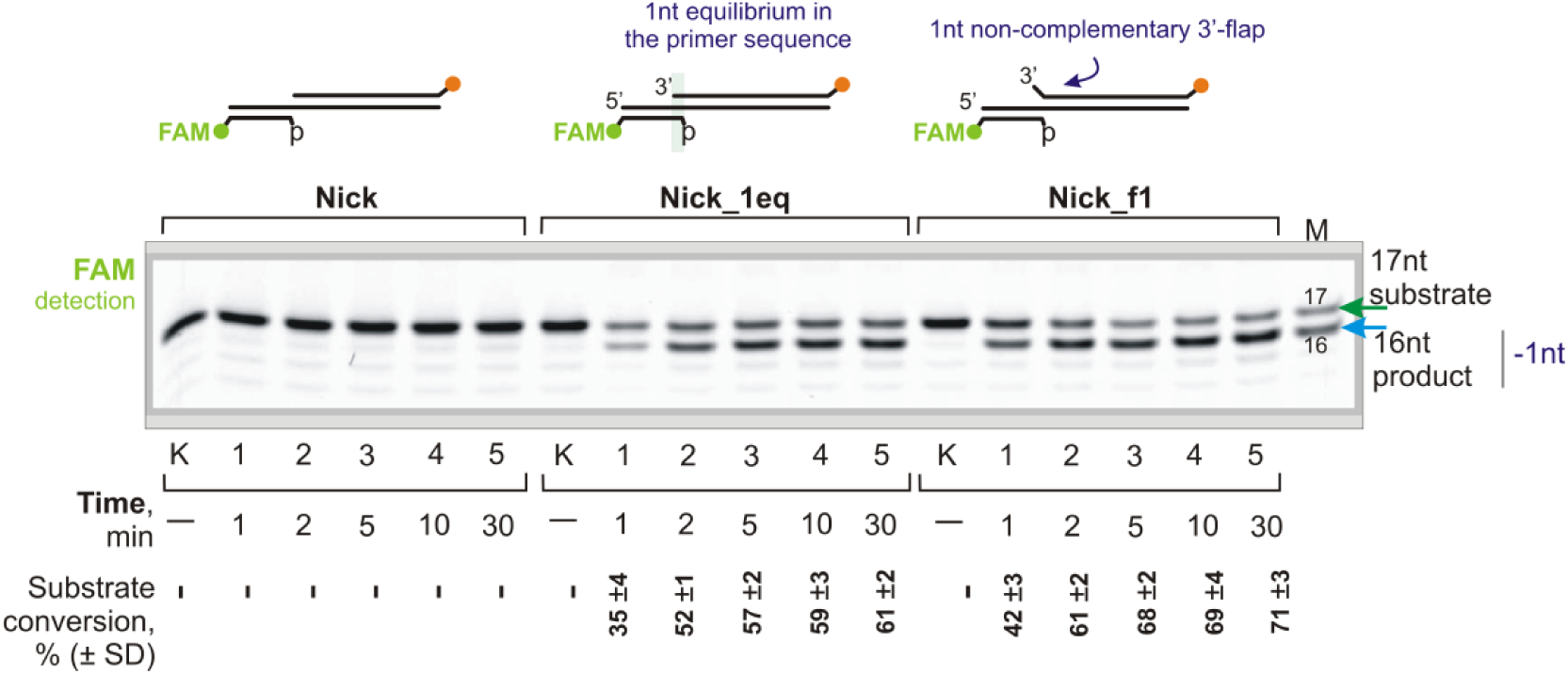
FEN1 is able to process non-complementary 3’-flap DNA (Nick_f1) better than complementary 3’-nucleotide DNA forming an equilibrium area (Nick_1eq). The names and schematic representations of the substrates are depicted above the gel; lime green and orange circles indicate FAM and TAMRA labels, respectively; the green arrow indicates the substrate DNA (flap-strand); the blue arrow indicates the 16nt cleavage product. Lane “k” is the DNA control; lane “M” is the length marker. The reaction mixtures contained 20 nM DNA and 20 nM FEN1.

### 3.5 FEN1 is activated by DNA-intermediates containing an equilibrium nucleotide area

It was previously found that the Polβ lyase rate constant is 20-fold higher than the polymerase rate constant [27]. Thus, Polβ rapidly removes the dRP-group generating a 1nt-gap and then much more slowly catalyzes the gap-filling reaction creating nicked DNA. Following this order of DNA-intermediate formation, we designed experiments to investigate exactly which DNA-structure “switches on” FEN1 catalytic activity (**Fig. 5A**). FEN1 did not catalyze gap1-DNA cleavage in the absence of dNTPs (lanes 1-3 in **Fig. 5A**) and remained catalytically inactive after gap1-to-nick conversion by Polβ in the presence of dTTP (lanes 4-7 in **Fig. 5A**). Indeed, during the normal SP-BER process, the final ligatable nick is not processed by FEN1 and is channeled directly to ligase [1], [3]. However, if Polβ inserted an additional nucleotide, generating a non-ligatable product (lane 4 for nick DNA), this DNA-intermediate was subsequently processed by FEN1, which restored the ligatable status (lanes 5-7 for nicked DNA). Overall, the equilibrium nucleotide area activates FEN1’s catalytic activity, enabling excision and conversion to a nick for passage to the subsequent ligation step.

**Fig. 5.**
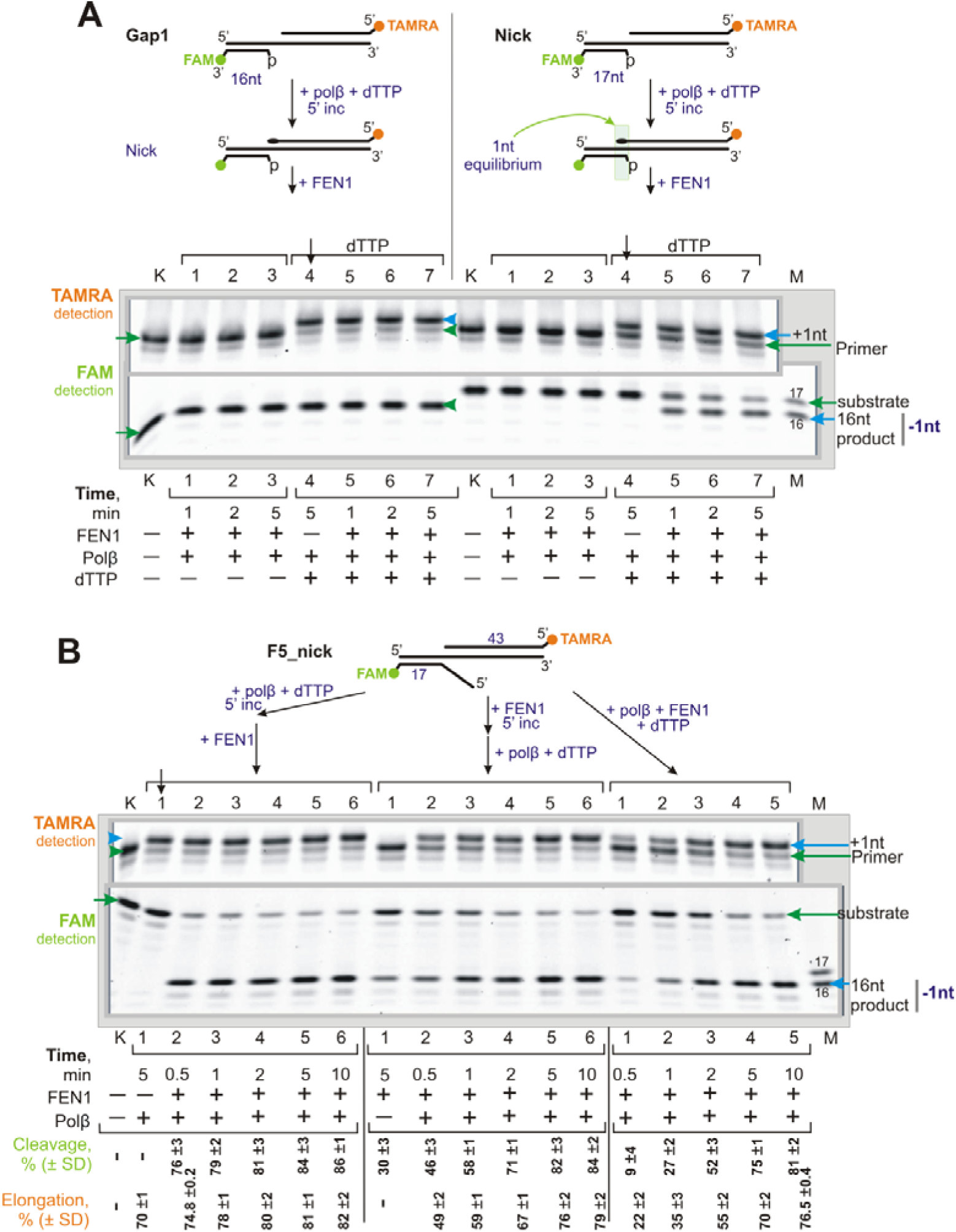
Polβ activates FEN1 cleavage activity by forming an equilibrium area. (A) After gap1 filling and conversion to a nick, FEN1 remains catalytically inactive. The subsequent formation of a 1nt equilibrium area leads to FEN1 activation. In the case of preformed nicked DNA, FEN1 is activated after the primer is elongated by 1nt, i.e., after the formation of a 1nt equilibrium area. (B) Different orders of incubation of FEN1 and Polβ with 5nt-flap DNA: Polβ is pre-incubated first, then FEN1 is added; FEN1 is pre-incubated first, then Polβ is added; or both proteins are added simultaneously. The yields of the cleavage and elongation reaction products are shown below the gel (%±SD).

Next, we performed kinetic experiments using DNA with a 5nt-flap and a nick, which mimicked the LP-BER DNA-intermediate **(Fig. 5B).** We varied the order of FEN1 and Polβ incubation with the substrate in the following scenarios: 1) Polβ was pre-incubated with DNA and dTTP prior to FEN1 being added; 2) FEN1 was incubated with DNA before (Polβ+dTTP) was added; or 3) FEN1, Polβ, and dTTP were added to the reaction mixture simultaneously. In the first scenario, Polβ inserted one nucleotide (lane 1) generating a 1nt equilibrium area, after which FEN1 processed almost the entire DNA within 30 seconds or less (lane 2). When FEN1 was pre-incubated, the product yield after 30 seconds of incubation with Polβ was approximately 50% (lane 2 in the second scenario). Adding both enzymes simultaneously resulted in a 50% product yield only after 2 min (lane 3, third scenario). Thus, even processing of the 5nt-long flap depends on the formation of an equilibrium area. Remarkably, in all three enzyme-addition scenarios, the DNA was completely converted to product within 10 minutes (lane 6 for all three scenarios).

The vertical black arrow denotes the lane with the Polβ elongation control before FEN1 supplementation; names and schematic representations of the substrates are shown above the gels; lime green and orange circles indicate FAM and TAMRA labels, respectively; green arrows indicate the position of the substrate DNA (primer or flap-strand); blue arrows indicate the elongation or cleavage products; the black oval denotes one incorporated dTMP. Lane “k” is the DNA control; lane “M” is the length marker. Reaction mixtures contained 20 nM DNA, 20 nM FEN1, 20 nM Polβ, and 1 µM dTTP (unless otherwise stated).

### 3.6 The position of the cleavage point is determined by the number of equilibrium nucleotides

Further, we expanded the equilibrium nucleotide area step-by-step to examine the shifting FEN1 cleavage points (**Fig. 6**). FEN1 does not catalyze the cleavage of nicked DNA (lanes k(F) and k(F+B)), but after the formation of a 1nt equilibrium area (lane 1 for nick), the product corresponding to the (-1) position is observed (lanes 2-4 for nick). After Polβ extends the primer by 2nt (lane 5 for nick), the cleavage position shifts to the (-2) position (lanes 6-8 for nick). Minor cleavage products at the (-1) and (-3) positions correspond to minor primer extension products of 1 and 3nt, respectively. Data obtained for DNA with a preformed 1nt equilibrium area (nick_1eq) are consistent with those for initially nicked DNA: when Polβ extends the primer by 1nt, forming a 2nt equilibrium area (lane 5 for nick_1eq), the flap-strand cleavage point shifts to the (-2) position (lanes 6-8 for nick-1eq). Similar results were obtained with F5_1eq DNA (**Fig. S7**): primer extension by 1nt drives a cleavage point to the next nucleotide in the duplex part.

**Fig. 6.**
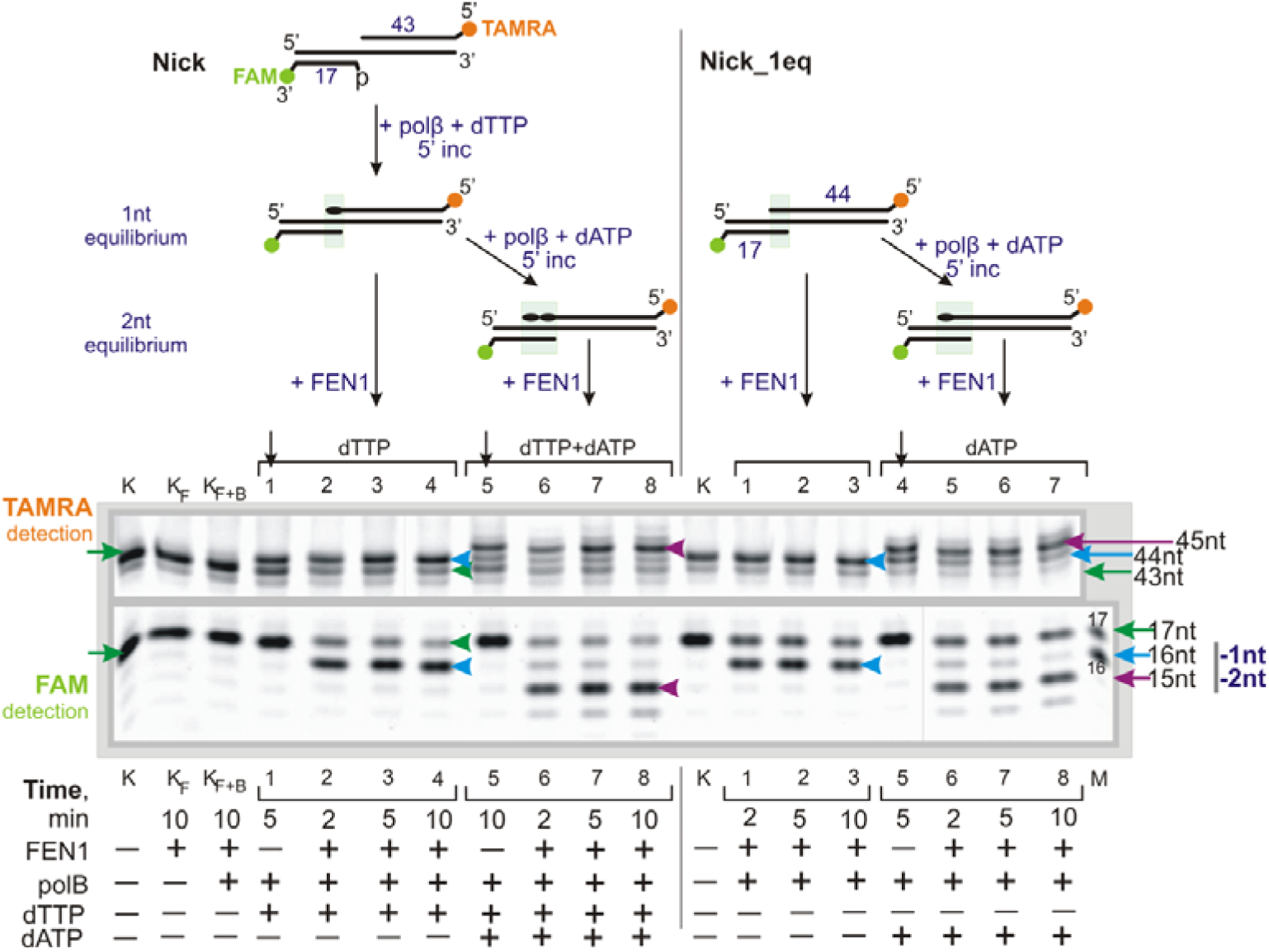
The number of equilibrium nucleotides determines the position of cleavage. Each step of primer extension shifts the cleavage point one nucleotide further into the duplex part from the previous point. The vertical black arrows denote the lane with the Polβ elongation control before FEN1 supplementation; names and schematic representations of the substrates are depicted above the gels; lime green and orange circles denote FAM and TAMRA labels, respectively; green arrows indicate the position of the substrate DNA (43nt primer or flap-strand); blue arrows point to the 44nt primer, the +1 elongation of the 43nt primer, or the (-1) cleavage product; plum arrows point to primer elongation products or (-2) cleavage products; the black oval denotes one incorporated dTMP or dAMP. Lane “k” is the DNA control; lane “M” - length marker. Reaction mixtures contained 20 nM DNA, 20 nM FEN1, 20 nM Polβ, and 1 µM dTTP and/or dATP (unless otherwise stated).

To clarify how the cleavage point depends on the number of equilibrium nucleotides, we performed experiments with DNA bearing a 5nt-flap combined with 0, 1, or 2 equilibrium nucleotide areas (see **Fig. S8**). The results showed that FEN1 can cleave DNA with or without a one-nucleotide equilibrium region at the (-1) position. A 2nt equilibrium region results in a (-2) cleavage position. We can therefore conclude that the cleavage point is located directly opposite the terminal equilibrium 3′-nucleotide of the primer, forming a potentially ligatable substrate. According to the structural data, FEN1 unpairs the 3′-terminal primer nucleotide and wedges it with the α2-α3 loop (see **Fig. S6C**). Notably, the FEN1 structure contains a specific loop bearing an “acidic block”, i.e., the EEGE residues 56–59, which block DNA from passing beyond the one-base pocket via charge repulsion (**Fig. S6B**) [17]. Thus, FEN1 binds to the terminal 3′-nucleotide independently of the total number of equilibrium nucleotides.

### 3.7 The FEN1 cleavage profile depends on dNTP concentration

The minimum patch size for LP-BER is reported to be 2nt, and at least half of repair events produce larger patches [28], [52]. We therefore hypothesize that longer products could be due to the initial lesion type or dNTP concentration at different stages of the cell cycle. Previous data suggest that the dNTP pool in S-phase increases from 1-5 µM up to 50 µM [49]. At the same time, the BER process operates throughout the cell cycle [1]. Therefore, we propose that long patches (and possibly corresponding long 5’-flaps) are generated at dNTP concentrations comparable to those in the S-phase.

The influence of dNTP concentration on the FEN1 cleavage point was estimated using DNA models of different BER intermediates, including gap1, nick (**Fig. 7**), as well as 5nt-flap, 1nt-flap with a nick, and a synthetic AP-site with a nick (**Fig. S8**). For nicked DNA at 1 µM dNTPs, Polβ extends primers by 1-2nt, generating a 1-2nt equilibrium area (**Fig. 7A**). After the equilibrium area forms, FEN1 cleaves the downstream primer at the (-1/-2) position creating a potentially ligatable product. The FEN1 cleavage product profile precisely mirrors the primer extension product profile. At 50 µM dNTPs, the primer extension product profile (+2, +3, +5, +6, and +8) corresponded to the cleavage product profile (-2, -3, -5, and -6).

**Fig. 7.**
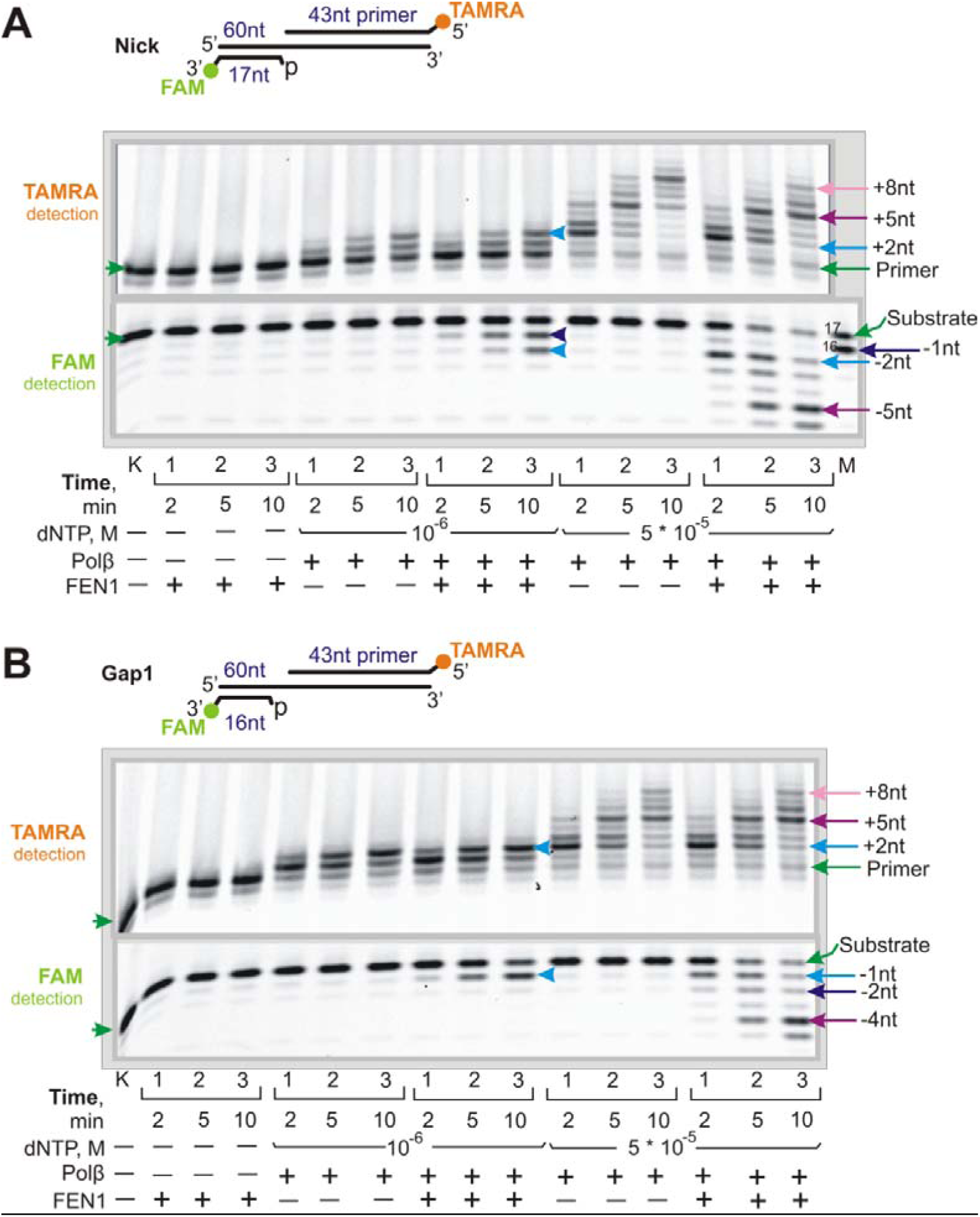
The profile of FEN1 cleavage products varies with Polβ activity at 1 and 50 µM dNTPs, using nicked DNA (**A**) and gap1 (**B**). The names and schematic representations of the substrates are shown above the gels; lime green and orange circles represent FAM and TAMRA labels, respectively; green arrows indicate the position of the substrate DNA (primer or flap-strand); light blue, dark blue and plum arrows indicate primer elongation and the corresponding cleavage products; lane “k” is the DNA control, and lane “M” is the length marker. Reaction mixtures contained 20 nM DNA, 20 nM FEN1, 20 nM Polβ, and dNTPs at the indicated concentrations.

An equivalent experiment was performed using a gap1 DNA (**Fig. 7B**). In this case, Polβ efficiently fills the gap and inserts a second nucleotide at 1 µM dNTPs, generating a 1nt equilibrium area. After equilibrium area formation, FEN1 cleaves the downstream primer at the (-1) position, generating a potentially ligatable product. With 50 µM dNTPs for 10 minutes, Polβ inserts up to 8nt, with pausing points at 2, 3, and 5nt. Notably, the number and intensity of FEN1 cleavage products (-1, -2, and -4) mirror the Polβ pausing points, shifted by 1nt, which is used for gap filling.

At 1 µM dNTPs, Polβ extends primer by 2nt with varying effectiveness. On gap1 DNA, Polβ inserts two nucleotides: the first fills the gap, and the second generates a 1nt equilibrium area. Interestingly, Polβ generates equilibrium nucleotide more effectively on gap1 DNA than on nicked DNA. This difference could be explained by Polβ’s ability to adopt a protein-DNA conformation when binding to the 5’-margin of the gap, which cannot be achieved with nicked DNA. Conversely, at 50 µM dNTPs, the primer extension product profiles (number and relative intensity) are identical for gap1 and nicked DNAs.

The same correspondence between the extension and cleavage products was obtained using F5_A_nick DNA containing a 1nt equilibrium area (see **Fig. S9C**). The cleavage product profile for this DNA (-1, -3, and -6) corresponded to the primer extension product profile, except for an additional 1nt shift (+2 and +5) due to the preformed 1nt equilibrium area, and the ability of FEN1 to cleave this DNA without support from Polβ. These experiments also demonstrate that Polβ extends the primer within the 5nt-flap DNA with efficiency comparable to that of gap1 DNA at 1 µM dNTPs, and with even greater efficiency at 50 µM dNTPs. We speculate that the five-nucleotide 5′-flap coupled with the equilibrium area induces more structural breathing, which may facilitate Polβ’s strand-displacement activity. Notably, the primer extension product profiles by Polβ at 50 µM dNTPs with and without FEN1 for gap1 and nick DNAs are identical. In the case of 5nt-flap DNA, FEN1 does not affect 2nt primer extension but causes an overall shortening of product length (see **Fig. S9C** at 50 µM dNTPs).

The cryo-EM structure of the FEN1-PCNA-DNA complex has recently been determined. In this structure, the DNA is endogenous and was co-purified from HEK293 cells transfected with a plasmid encoding PCNA [19]. This endogenous DNA is located in the catalytic center of FEN1 and contains at least a two-nucleotide 5’-fragment (it may contain more, but the rest of the 5’-fragment is not traceable) and a one-nucleotide 3’-fragment, forming an equilibrium zone (**Fig. S6C**). Combining these structural data with our biochemical results, we conclude that DNA containing an extended equilibrium area is a “native substrate” for FEN1.

### 3.8 The DNA substrate is channeled from Polβ to FEN1 within the Polβ-FEN1-DNA ternary complex

To determine whether FEN1 joins the Polβ-DNA complex or Polβ dissociates to allow FEN1 binding, we performed EMSA experiments with the FEN1 substrates: pTHFp_1eq and F5_1eq, as well as with a nicked DNA modeling the cleavage reaction product (**Fig. 8A**). We observed ternary complex formation with all three DNAs. These experiments were conducted under non-catalytic conditions in the presence of EDTA, since metal ions do not necessarily need to be present in the active site for the initial FEN1 binding and DNA bending [47]. When we excluded EDTA from the reaction mixture, the ternary complex was accompanied by products of the cleavage reaction (lanes 9-13 in **Fig. S10A** and lanes 6-10 in **Fig. S10C**). To examine the ability of FEN1 to cleave DNA without the addition of Mg^2+^/Mn^2+^ ions, we used two DNA substrates comprising a 5nt-flap in combination with a nick or a 1nt equilibrium region (**Fig. S10B**). Indeed, this experiment confirms a previous finding that FEN1 is able to cleave F5_1eq DNA without the addition of Mg^2+^/Mn^2+^ (lane 1) possibly because these metal ions are captured by the protein during purification. Notably, the addition of dNTPs does not affect the efficiency of ternary complex formation (lanes 6-8 at **Fig. S10A**), although dNTP binding to the Polβ-DNA binary complex is known to result in a more closed conformation of Polβ [53]. Interestingly, the ternary complex was detected even after the FEN1 catalysis reaction was completed and the DNA was converted to the nicked structure (**Fig. 8A, Fig. S10A,** and **C**).

**Fig. 8.**
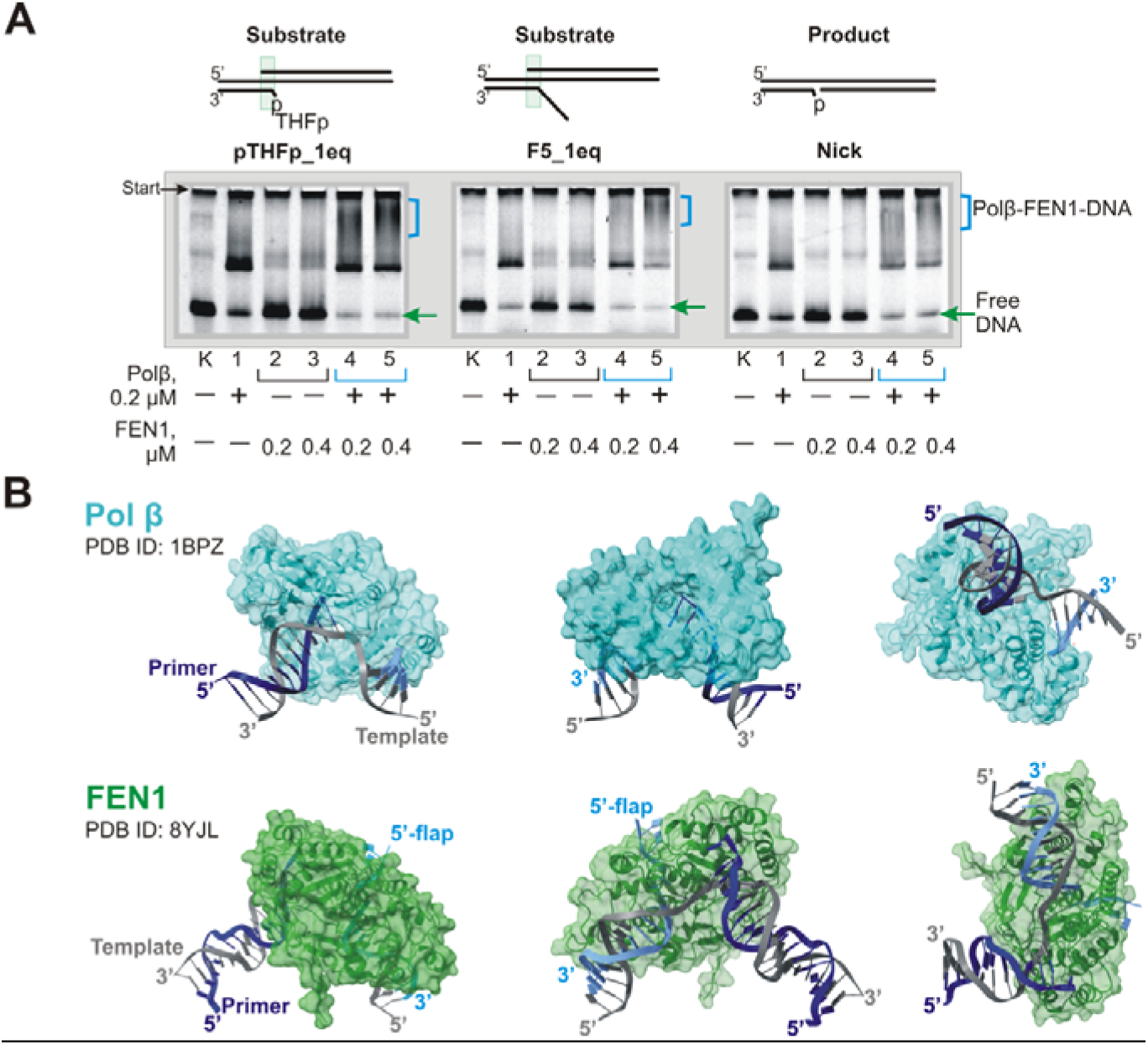
(**A**) The Polβ-FEN1-DNA ternary complex was detected with FEN1’s substrate DNAs (pTHFp_1eq and F5_1eq) and product nicked DNA. A schematic representation of the substrates is shown at the top; the green arrow indicates the position of free, unbound DNA; blue staples indicate putative Polβ-FEN1-DNA complexes. Lane “k” is the DNA control. Reaction mixtures (10 μL) contained 20 nM DNA, 200 nM Polβ, 5 mM EDTA, and FEN1 at a concentration of 200 or 400 nM. (**B**) Binding of Polβ (cyan) does not interfere with binding of FEN1 (green) to the downstream duplex part region. Polβ binds predominantly to the upstream duplex part of the nicked DNA (X-ray structure, PDB ID: 1BPZ). FEN1 initially binds to the downstream duplex part containing the flap-strand (cryo-EM data of FEN1 in complex with product 5’-flap DNA, PDB ID: 8YJL). Following the initial FEN1 binding to the downstream duplex part, this is likely further accompanied by internal architectural perturbations within the Polβ-FEN1-DNA complex to hand off the DNA-substrate to FEN1. Figure preparation was conducted using UCSF ChimeraX.

According to the structural data (**Fig. 8B**), the largest Polβ-DNA interface is formed by the primer-side duplex region (X-ray structure of the Polβ-nick-DNA complex, PDB ID: 1BPZ). Thus, the downstream duplex part remains exposed for the initial FEN1 binding. Additionally, Polβ binds both gapped and nicked DNA with a 90° kink occurring precisely at the 5’-phosphodiester linkage of the templating residue [54]. FEN1 recognizes the bent DNA itself, owing to its intrinsic ability to recognize DNA with a sharp bend flanked by duplex portions on either side [17], or recognizes the whole Polβ-DNA complex via direct Polβ-FEN1 interaction [23], [24]. FEN1 then joins the Polβ-DNA complex by binding to the downstream duplex part. In the next step FEN1 has to gain access to the 5’-flap-strand and to the 3’-nucleotide of the primer, which are buried within the Polβ complex. Therefore, FEN1 must either displace Polβ, or some architectural perturbation occurs within the Polβ-FEN1-DNA complex to handoff the DNA-substrate to FEN1. Given that the ternary complex forms with nicked DNA, it can be assumed that Polβ does not dissociate from the complex, and that DNA handoff occurs within it.

## 4. Discussion

The current model of BER proposes that the repair process utilizes the long-patch pathway when the AP-site is either oxidized or reduced, such that Polβ cannot eliminate the modified sugar via lyase activity. Without the formation of a catalytically competent complex with the 5’-margin of the gap, Polβ starts to perform strand-displacement synthesis. The ability of Polβ to “test” the 5’-side of a gap can be employed to scan for the presence of DNA damage [55]. Our *in vitro* experiments demonstrate that Polβ can initiate strand-displacement synthesis when the 5’-margin contains either a tetrahydrofuran or a normal 5’-phosphate in gap or nicked DNA [56], [57], [58]. The subsequent equilibrium nucleotide area is recognized by FEN1, in particular which probe the 3’-nucleotide for the presence of a 3’-OH group. The obtained results provide the opportunity to assimilate our biochemical findings together with those of previous studies [35], [59], with existing structural data in order to reveal the LP-BER substrate probing mechanism: 1 – Polβ checks the 5’-side and catalyzes extension of the 3’-side; 2 – FEN1 tests the 3’-side nucleotide for ligatable status and catalyzes cleavage of the 5’-side. The subsequent elimination of the equilibrium area leads to restoration of the ligatable structure. Therefore, this mechanism, in contrast to “gap-translation”, excludes further strand-displacement synthesis. In addition, strand-displacement synthesis in the BER can be inhibited by regulatory proteins – poly(ADP-ribose)polymerases 1 and 2 (PARP1 and 2) [58], [60]. On the other hand, PARP1 and 2 and the poly(ADP-ribose) they synthesized, may serve as triggers in the formation of cellular compartments that facilitate the BER process [61].

The structural specificity of FEN1 is provided by the organization of structurally identical enzyme-substrate complexes, regardless of the overall length of the equilibrium nucleotide region in the DNA substrate. FEN1 binds to the 3’-primer nucleotide and forms a double-flap branching structure comprising a 3’-flap of one nucleotide and the corresponding 5’-flap. Subsequently, FEN1 cleaves the bound DNA intermediate at the (-1) position from the branch point into the duplex part of the flap-strand (**Fig. S11**).

Interestingly, these data raise the question of what repair events could occur after Polβ inserts a non-complementary nucleotide and FEN1 recognizes the 3’-flap and cleaves the downstream primer at the (-1) position, thereby generating a pseudo-gap with a 3’-flap DNA.

Such mutagenic DNA intermediates might be repaired by the recently discovered LP-BER sub-pathway, which involves XPF-ERCC1 cleavage of a 3’-flap [62].

Due to the simultaneous detection of the extension and cleavage products, we found that these products are symmetrical. This fact allows us to suggest that when Polβ pauses, FEN1 joins the Polβ-DNA complex and cleaves the flap. Thereafter, the DNA sequence could facilitate the start of LP-BER start and influence the repair patch length. Indeed, *Xue et al.* have recently reported that the surrounding DNA sequence influences LP-BER distribution; specifically, changing the downstream sequence to be less G-rich gave rise to longer repair patches [29]. Additionally, our data demonstrate that an equilibrium nucleotide area longer than 2nt can be eliminated by means of a single FEN1 cleavage event, rather than through several one-by-one steps.

Direct protein-protein interaction between FEN1 and Polβ has been demonstrated previously [23], [24]. Although the ternary Polβ-FEN1-DNA complex was not detected before with the use of EMSA [26], [27], this complex was demonstrated to form through crosslinking of FEN1 and Polβ to the central BER intermediate, followed by their co-immunoprecipitation [25]. Without direct handover from Polβ to FEN1, that is, if Polβ dissociates before FEN1 binds, the DNA-intermediate is left unprotected and vulnerable to harmful nuclease activities, recombination events, or cell-death signaling. Therefore, it seems that there should be a mechanism that channels the DNA intermediate from Polβ to FEN1, avoiding this potential “unprotected exposure”. Indeed, EMSA experiments clearly show that a ternary Polβ-FEN1-DNA complex can be formed with all of FEN1’s substrate DNAs. Moreover, these proteins remain in the complex even after the FEN1 catalytic reaction is completed and the DNA is converted to the nicked structure. Thus, we cannot exclude the possibility that FEN1 does not dissociate from the Polβ-DNA complex during the subsequent strand-displacement synthesis. These data therefore suggest a rapid DNA handover through architectural rearrangements within the Polβ-FEN1-DNA complex.

During the maturation of Okazaki fragments, the strand-displacement activity of Polδ and the 5’-flap cleavage activity of FEN1 act processively through the iterative 1-3nt steps of nick translation, ultimately removing the RNA/DNA primer [63]. However, the details of the handover mechanism inside the Polδ-FEN1-DNA-PCNA complex are unknown, while the rates of the nick translation imply that Polδ and FEN1 actively hand off their products [37].

According to our data, FEN1 cleaves the downstream strand directly opposite the terminal equilibrium 3’-nucleotide, generating a product that can potentially be ligated.

Therefore, the size of repair patches depends on the total number of equilibrium nucleotides produced by Polβ activity, particularly by synthesis pausing. The repair patches determined for 1 µM dNTPs, corresponding to the dNTP concentration throughout the cell cycle except S-phase, showed 2nt products predomination. That is consistent with recent HEK293 cell extract data where the 2nt product dominated for different repair substrates [29]. Polβ activity increases as dNTP concentration rises towards its S-phase level. We hypothesize that LP-BER repair patches become longer when the repair process occurs during S-phase, with its elevated dNTP concentration, and shorter when the dNTP concentration is reduced to basal levels (see **Fig. 9**).

**Fig. 9.**
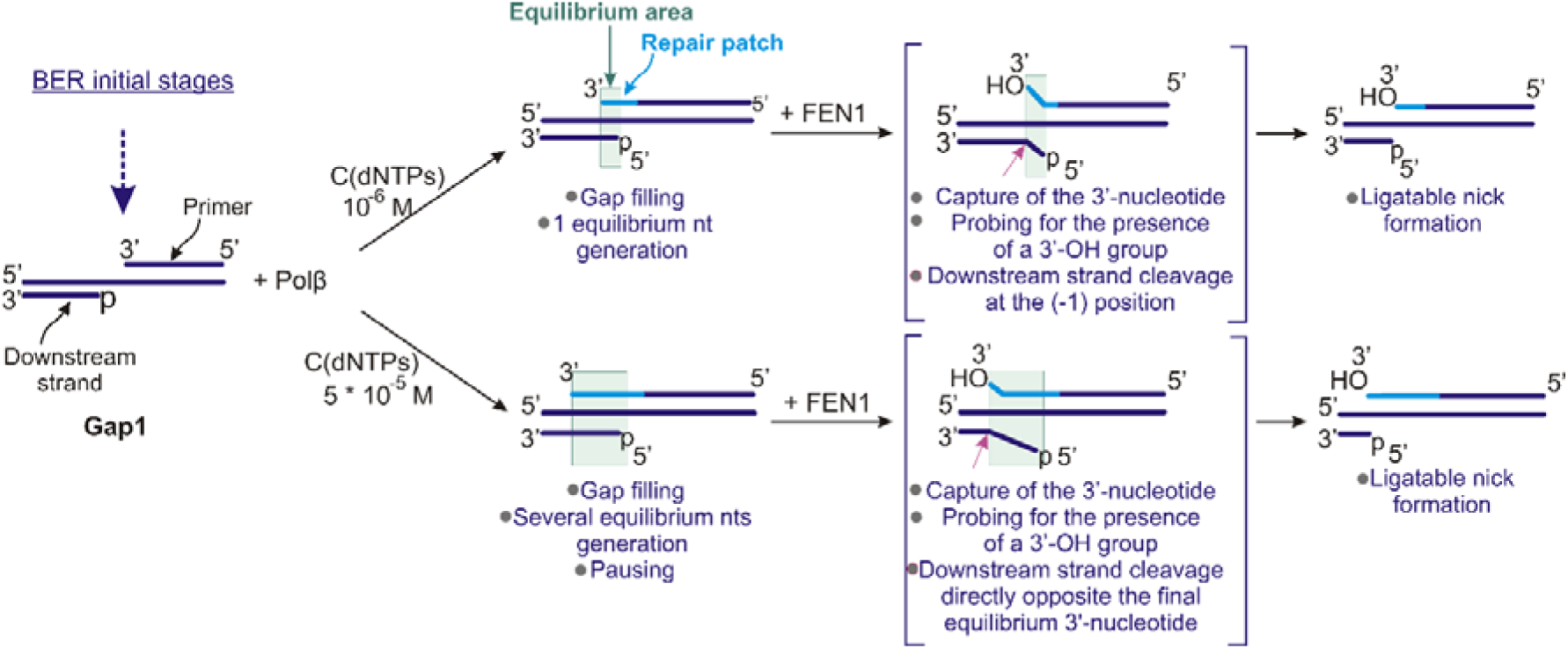
A model for the formation of DNA intermediates that “switch on” FEN1 catalytic activity and the dependence of repair patch size on dNTP concentration. In case of gap1 DNA and 1 µM dNTPs, Polβ efficiently fills the gap and inserts a second nucleotide, creating a 1nt equilibrium area. After formation of the equilibrium area, FEN1 cleaves the downstream primer at the (-1) position, forming a product that can potentially be ligated. Increasing the dNTP concentration to 50 µM leads to the primer extension by several nucleotides, accompanied by multiple pausing points. After Polβ pauses, FEN1 joins the Polβ-DNA complex to cleave the flap. The number and intensity of FEN1 cleavage products mirror the Polβ pausing points, with a 1nt shift, which is used to fill the initial gap. The cleavage point is located directly opposite the terminal equilibrium 3’-nucleotide of the primer, and the resulting product is suitable for ligation.

During replication, DNA binding factors or nucleosome obstacles may cause Polδ pausing/stalling, thereby promoting subsequent DNA handover from FEN1 to Lig1 [64]. Acute depletion of Lig1 in yeast leads to additional nick translation [65]. It is noteworthy that our model experiments do not include DNA ligases, which are responsible for nick sealing in the BER process. In the presence of ligase, DNA may be channeled from FEN1 to Lig1 rather than back to Polβ, thus suggesting that DNA ligase is responsible for limiting the length of the repair patch [66]. Additionally, channeling of nicked DNA from FEN1 to Lig1 may require an “unbent” DNA conformation, whereas handover back to Polβ could instead be performed at a bent conformation corresponding to the FEN1 catalytic complex [51], [64].

## 5. Conclusion

Taken together, our biochemical analyses reveal the mechanism of formation of the DNA intermediate that activates the FEN1 catalytic reaction. Our results indicate that the FEN1-Polβ interplay occurs within the Polβ-FEN1-DNA complex through steps of generation-elimination of equilibrium nucleotide areas, rather than through a “gap translation” mechanism. Following initial binding to the nascent equilibrium nucleotide site, FEN1 bends the DNA intermediate and captures the 3’-nucleotide of the primer. The presence of a 3’-OH group on the 3’-nucleotide of the primer is necessary for catalytic activation. Owing to FEN1’s structural ability to organize a catalytically active FEN1-DNA complex based on the 3’-nucleotide, the resulting complex forms identically with DNA of varying flap lengths. The 5’-flap within this complex is cleaved precisely at the (-1) position from the branch point within the duplex part. The resulting cleavage product is a ligatable nicked DNA.

## CRediT authorship contribution statement

**Yuliya S. Krasikova**: Investigation, Methodology, Validation, Writing - Original Draft, Visualization; **Ekaterina A. Maltseva**: Polβ purification, Validation, Writing - Review & Editing; **Svetlana N. Khodyreva**: FEN1 purification, Writing - Review & Editing; **Nadejda I. Rechkunova**: Validation, Writing - Review & Editing, Project administration; **Olga I. Lavrik**: Conceptualization, Supervision, Writing - Review & Editing, Funding acquisition.

## Data availability

No data was used for the research described in the article.

