## Supplementary Materials for "Interplay between flap endonuclease 1 and DNA polymerase β in long patch base excision repair: new facets in mechanism"

**Table S1.** Oligonucleotides used in this study for preparing DNA substrates. Flap-forming sequences are underlined. The fluorescent labels FAM and TAMRA are located at the indicated positions.

| **Name** | **Sequences 5’ → 3’** |
| --- | --- |
| **60** | 5’ –  Ctatggcgaggcgattatcaacccatttagtcgtaatagtgaagagtcacgacaacatcg |
| **16pF** | 5’ – **p**AATCgCCTCgCCATAg – 3’ – **FAM** |
| **17pF** | 5’ – **p**TAATCgCCTCgCCATAg – 3’ – **FAM** |
| **pTHFpF** | 5’ – **pTHFp**TAATCgCCTCgCCATAg – 3’ – **FAM** |
| **18pF**  **18F** | 5’ – **p**T TAATCgCCTCgCCATAg – 3’ – **FAM**  5’ – T TAATCgCCTCgCCATAg – 3’ – **FAM** |
| **19F** | 5’ – CT TAATCgCCTCgCCATAg – 3’ – **FAM** |
| **20F** | 5’ – ACT TAATCgCCTCgCCATAg – 3’ – **FAM** |
| **22F**  **22AF** | 5’ – CAACT TAATCgCCTCgCCATAg – 3’ – **FAM**  5’ – CAACA TAATCgCCTCgCCATAg – 3’ – **FAM** |
| **43up**  **43upT** | 5’ – Cgatgttgtcgtgactcttcactattacgactaaatgggttga – 3’  5’ – Cga[dt-**TAMRA**]gttgtcgtgactcttcactattacgactaaatgggttga–3’ |
| **44up_f1** | 5’ – Cga[dt-**TAMRA**]gttgtcgtgactcttcactattacgactaaatgggttgac |
| **44upT** | 5’ – Cga[dt-**TAMRA**]gttgtcgtgactcttcactattacgactaaatgggttgat |

**Table S2.** DNA substrates used in this study. The template 60mer oligonucleotide (5’-3’ view) is underlined. The flap-strand is shown in purple. The fluorescent labels FAM and TAMRA are designated as **F** and **T**, respectively. Equilibrium areas are highlighted in light blue.

| Substrate name | Substrate sequence |
| --- | --- |
| **Gap1**  (60+16pF+43upT) | agttgggtaaatcagcattatcacttctcagtgctgttgtagc**T**  Ctatggcgaggcgattatcaacccatttagtcgtaatagtgaagagtcacgacaacatcg  **F**gataccgctccgctaap |
| **Nick**  (60+17pF+43up) | agttgggtaaatcagcattatcacttctcagtgctgttgtagc  Ctatggcgaggcgattatcaacccatttagtcgtaatagtgaagagtcacgacaacatcg  **F**gataccgctccgctaatp |
| **Nick_f1**  (60+17pF+44up_f1) | cagttgggtaaatcagcattatcacttctcagtgctgttgtagc**T**  Ctatggcgaggcgattatcaacccatttagtcgtaatagtgaagagtcacgacaacatcg  **F**gataccgctccgctaatp |
| **Nick_1eq**  (60+17pF+44upT) | tagttgggtaaatcagcattatcacttctcagtgctgttgtagc**T**  Ctatggcgaggcgattatcaacccatttagtcgtaatagtgaagagtcacgacaacatcg  **F**gataccgctccgctaatp |
| **pTHFp_nick**  **(**60+pTHFpF+43up) | agttgggtaaatcagcattatcacttctcagtgctgttgtagc  Ctatggcgaggcgattatcaacccatttagtcgtaatagtgaagagtcacgacaacatcg  **F**gataccgctccgctaat  pTHFp |
| **F1p_nick**  **(**60+18pF+43up) | agttgggtaaatcagcattatcacttctcagtgctgttgtagc  Ctatggcgaggcgattatcaacccatttagtcgtaatagtgaagagtcacgacaacatcg  **F**gataccgctccgctaat  tp |
| **F1_nick**  **(**60+18F+43up) | agttgggtaaatcagcattatcacttctcagtgctgttgtagc  Ctatggcgaggcgattatcaacccatttagtcgtaatagtgaagagtcacgacaacatcg  **F**gataccgctccgctaat  t |
| **F2_nick**  **(**60+19F+43up) | agttgggtaaatcagcattatcacttctcagtgctgttgtagc  Ctatggcgaggcgattatcaacccatttagtcgtaatagtgaagagtcacgacaacatcg  **F**gataccgctccgctaat  tc |
| **F3_nick**  **(**60+20F+43upT) | agttgggtaaatcagcattatcacttctcagtgctgttgtagc  Ctatggcgaggcgattatcaacccatttagtcgtaatagtgaagagtcacgacaacatcg  **F**gataccgctccgctaat  tca |
| **F5_nick**  **(**60+22F+43upT) | agttgggtaaatcagcattatcacttctcagtgctgttgtagc**T**  Ctatggcgaggcgattatcaacccatttagtcgtaatagtgaagagtcacgacaacatcg  **F**gataccgctccgctaat  tcaac |
| **F5_1eq**  **(**60+22F+44upT) | tagttgggtaaatcagcattatcacttctcagtgctgttgtagc**T**  Ctatggcgaggcgattatcaacccatttagtcgtaatagtgaagagtcacgacaacatcg  **F**gataccgctccgctaat  tcaac |
| **F5_A_1eq**  **(**60+22AF+44upT) | agttgggtaaatcagcattatcacttctcagtgctgttgtagc**T**  Ctatggcgaggcgattatcaacccatttagtcgtaatagtgaagagtcacgacaacatcg  **F**gataccgctccgctaat  acaac |
| **F5_A_2eq**  **(**60+22AF+44upT) | tagttgggtaaatcagcattatcacttctcagtgctgttgtagc**T**  Ctatggcgaggcgattatcaacccatttagtcgtaatagtgaagagtcacgacaacatcg  **F**gataccgctccgctaat  acaac |

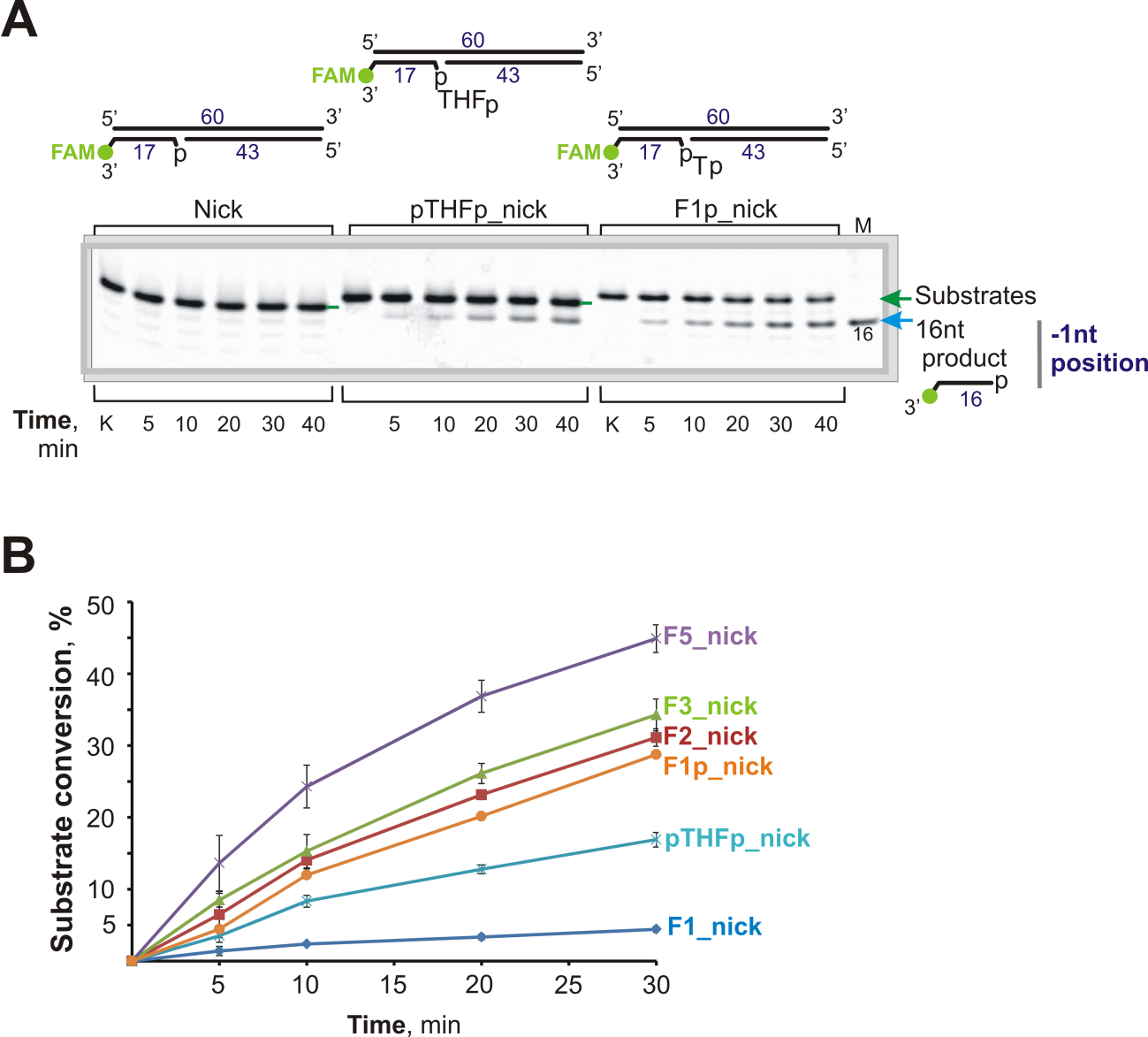

**Figure S1.** (**A**) FEN1 cleaves the 1nt-flap DNA containing a phosphate (F1p_nick) and DNA containing a synthetic analogue of the AP-site (tetrahydrofuran, THF) with a phosphate (pTHFp_nick) at the 5′-margin, at the same position. A schematic representation of the substrates is shown at the top; the lime green circle indicates the FAM-label position; the green arrow indicates the position of the substrate DNA; the blue arrow indicates the 16nt product DNA. Lane “k” is the DNA control; lane “M” is the length marker. Reaction mixtures contained 20 nM DNA and 20 nM FEN1. (**B**) The percentage of substrate conversion indicates the conversion of substrate to product. The product yield (±SD) for each substrate was calculated based on the experiments presented in panel (**A**) and Fig. 1.

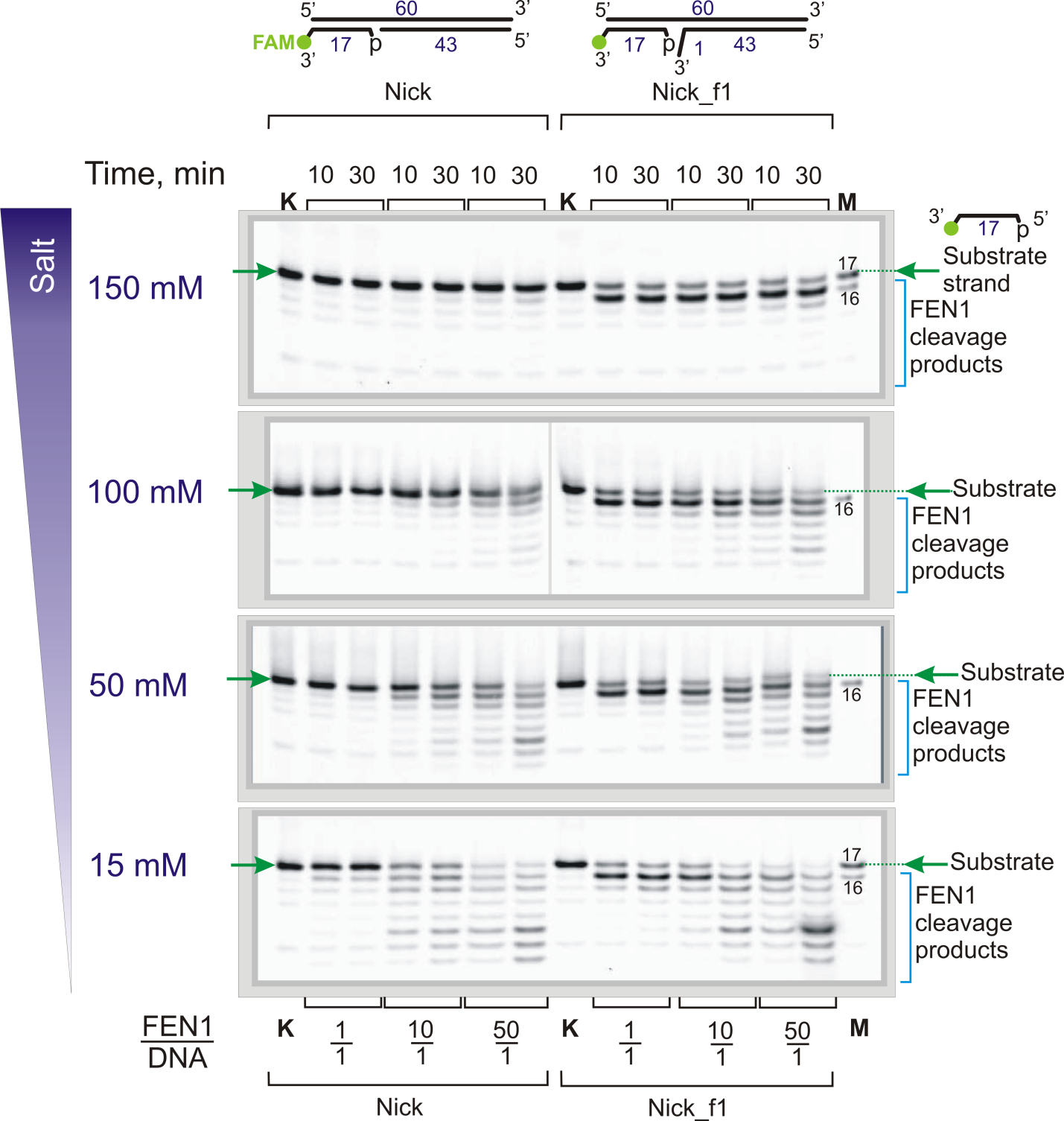

**Figure S2.** The exonuclease activity of FEN1 depends on the concentration of monovalent metal ions. At physiological concentrations of monovalent metal ions (100–150 mM), FEN1 is incapable of cleaving Nick DNA. However, it can cleave a Nick with a 3′ non-complementary flap directly at the (-1) position within the duplex part (the resulting product is 16nt). Decreasing the salt concentration results in the appearance of exonuclease activity and step-by-step degradation of the downstream primer. A schematic representation of the substrates is depicted at the top; the lime green circle indicates the FAM-label position; green arrows indicate the position of the substrate DNA; blue staples indicate the cleavage products. Lane “k” is the DNA control; lane “M” is the length marker. Reaction mixtures contained 20 nM DNA and FEN1 at the indicated enzyme/DNA ratio.

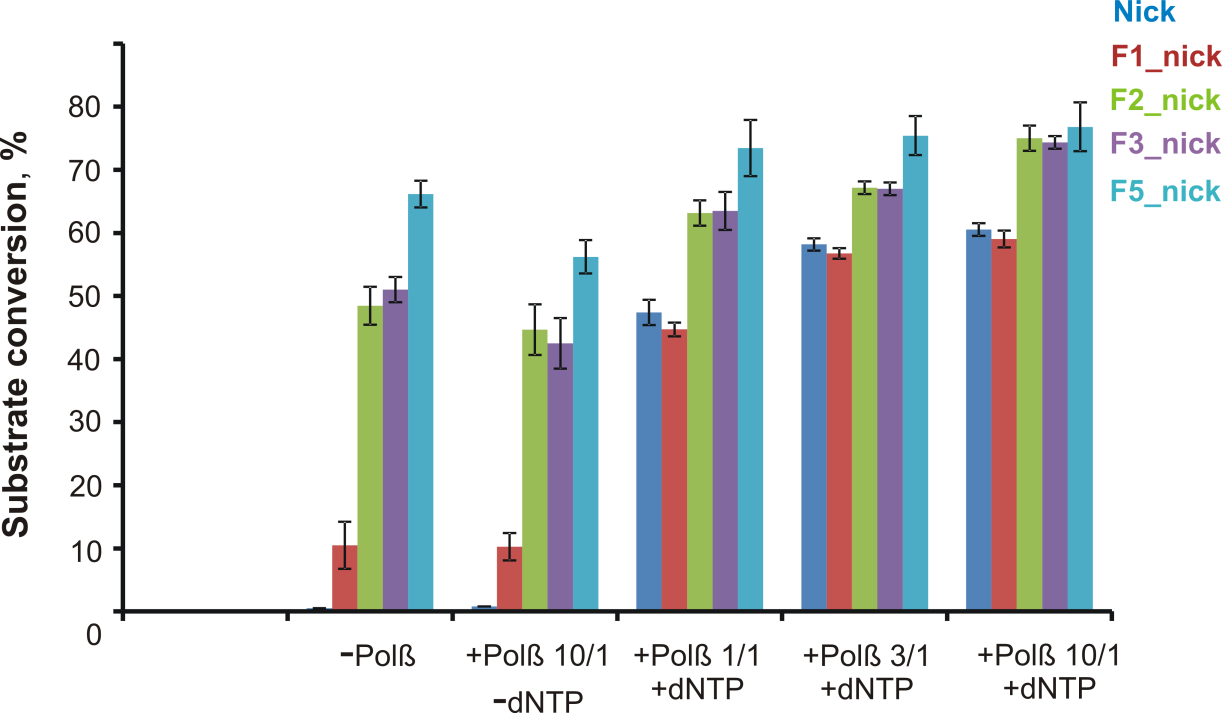

**Figure S3.** Polβ increases the efficiency of DNA cleavage by FEN1 in the presence of 1 µM dNTPs.

Polβ significantly increases the yield of cleavage products for nick and F1_nick DNAs. The cleavage efficiency of 2-, 3-, and 5nt-flap DNAs increased moderately. The percentage of substrate conversion indicates the conversion of substrate to product. The names of the DNA substrates and their corresponding color codes are presented on the right. The product yield (±SD) for each substrate was calculated based on the experiments presented in Fig. 2A. Reaction mixtures contained 20 nM DNA, 20 nM FEN1, 1 µM dNTPs, and Polβ at the indicated enzyme-to-DNA ratio.

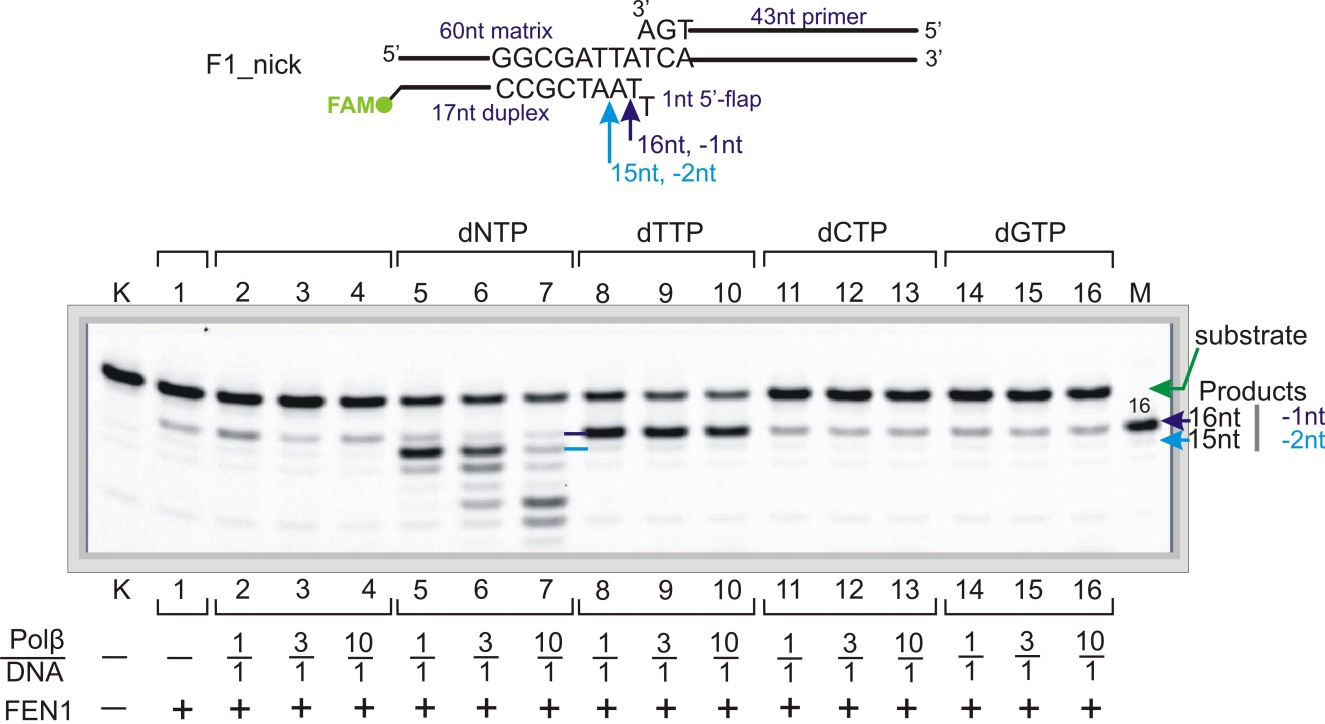

**Figure S4.** Polβ extends the primer in 1nt-flap DNA, providing FEN1-catalyzed DNA cleavage. The DNA contains the FAM-label at the 3′-end of the downstream primer, as shown in the schematic above the gel. The name and schematic representation of the substrate are depicted above the gel; the lime green circle denotes FAM-label; the green arrow indicates the position of the substrate DNA (flap-strand); the dark blue and light blue arrows/lines indicate 16nt and 15nt-length cleavage products, respectively. Lane “k” is the DNA control; lane “M” is the length marker. Reaction mixtures contained 20 nM DNA, 20 nM FEN1, Polβ at the indicated enzyme-to-DNA ratio, and 1 µM dNTPs (all four or individually).

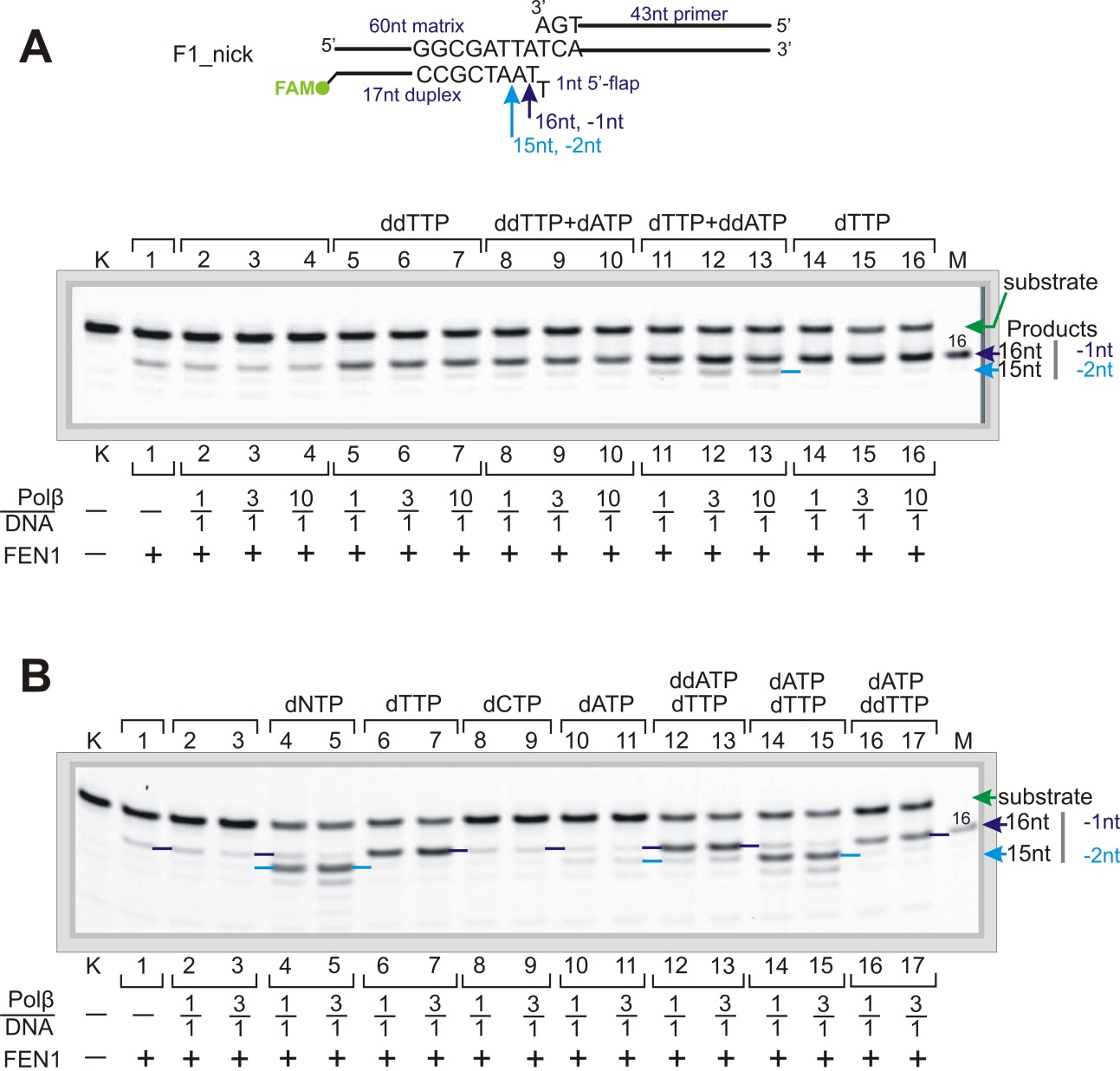

**Figure S5.** The cleavage stimulation effect depends on the presence of the 3ʹ-OH group and is therefore reduced by the incorporation of dideoxynucleotide (**A** and **B**). The name and schematic representation of the substrate are shown above the gels; the lime green circle denotes the FAM-label; the green arrow indicates the position of the substrate DNA (flap-strand); dark blue and light blue arrows/lines indicate 16nt and 15nt-length cleavage products, respectively. Lane “k” is the DNA control; lane “M” is the length marker. The reaction mixtures contained 20 nM DNA, 20 nM FEN1, Polβ at the indicated enzyme/DNA ratio, and 1 µM dNTPs or ddNTPs (all four, individually or in the indicated combination).

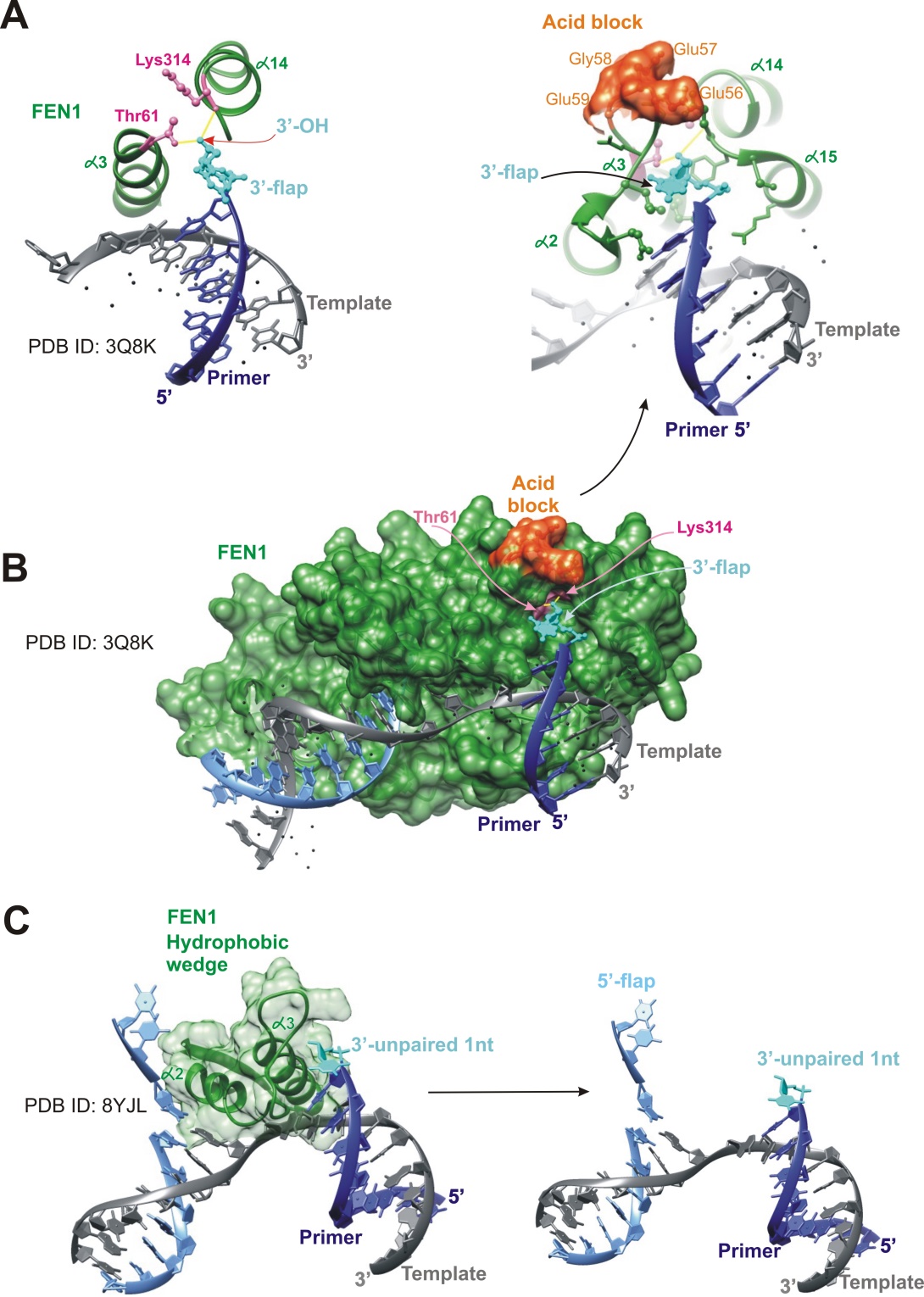

**Figure S6. Structural data on FEN1 contacts with the 3’-nucleotide:**

**A.**	Crystal structure data of FEN1 binding to the 3’-OH group, PDB ID: 3Q8K [Tsutakawa SE et al., Cell, 2011]. The Lys314 backbone carbonyl from the α14-helix and the Thr61 hydroxyl from the α3-helix form hydrogen bonds with the 3′-OH group. The FEN1 helices are shown in green, the defined aa are labeled in pink, the 3’-nucleotide is colored light blue, and the yellow lines correspond to the hydrogen bonds.

**B.**	The 3’-nucleotide binds within the 3’-nucleotide binding pocket, PDB ID: 3Q8K. The FEN1 structure contains a specific loop with an ‘acidic block’ (colored red), i.e., the EEGE residues 56-59, which block DNA from passing beyond the one-base pocket via charge repulsion. The color scheme is identical to that in (**A**).

**C.**	From the X-ray and cryo-EM data, PDB ID: 8YJL, we know that the α2-α3 hydrophobic loop unpairs the nearest 3’-primer base pair and stabilizes the forming 3’-flap. Endogenous DNA co-purified from HEK293 cells with the FEN1-PCNA complex contains at least a 2nt-long 5’-flap (it may be longer, but the rest of the 5′-flap is untraceable) and a 1nt 3’-flap that forms an equilibrium area [Tian Y et al., EMBO J., 2025].

Figure preparation was conducted using UCSF Chimera.

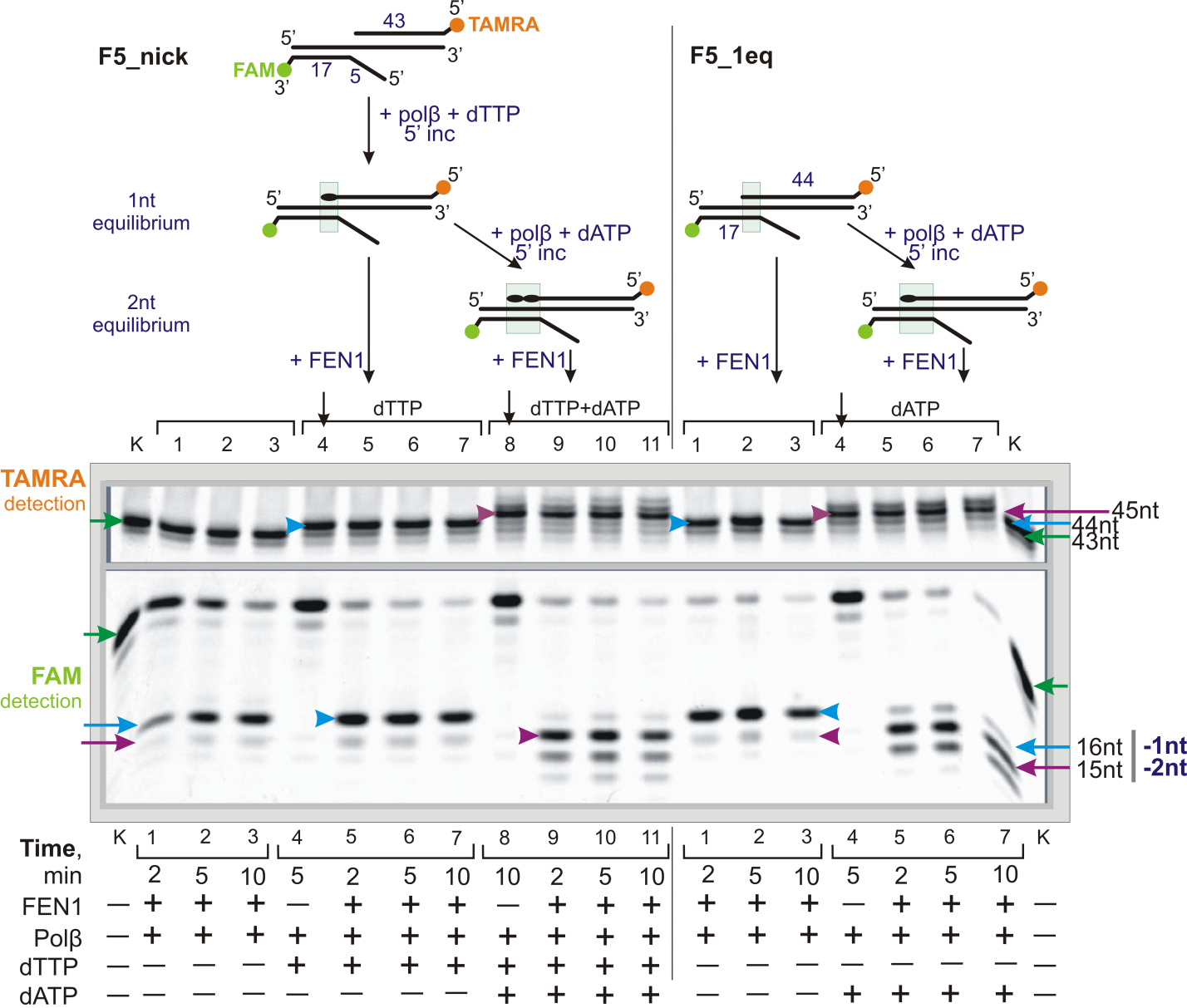

**Figure S7.** The location of the cleavage point depends on the number of nucleotides incorporated into the primer of the DNA containing the 5nt-flap. When Polβ forms a 1nt equilibrium area, the cleavage point is located at the -1nt position (a 16nt product). If a pre-formed 1nt equilibrium area is present, FEN1 cleaves the flap-strand at the same position. Subsequent primer extension generates a 2nt equilibrium nucleotide area, thereby shifting the cleavage point to the (-2nt) position (a 15nt product).

In the schematic representation of the reaction: the initial names and substrates are depicted at the top; the lime green and orange circles indicate FAM and TAMRA labels, respectively; the light green rectangle indicates the equilibrium nucleotides area; the black oval indicates one incorporated dTMP or dAMP. In the gels: the vertical black arrows in lanes 4 and 8 denote the Polβ elongation controls before the supplementation of FEN1; the green arrows indicate the position of the substrate DNA (43nt-primer or flap-strand); the blue arrows indicate the 44nt-primer, the +1 elongation of the 43nt-primer, or the (-1) cleavage product; the plum arrows indicate the primer elongation products or the (-2) cleavage products. Lane “k” is the DNA control. Reaction mixtures contained 20 nM DNA, 20 nM FEN1, 20 nM Polβ, and 1 µM dTTP or dATP (unless otherwise stated).

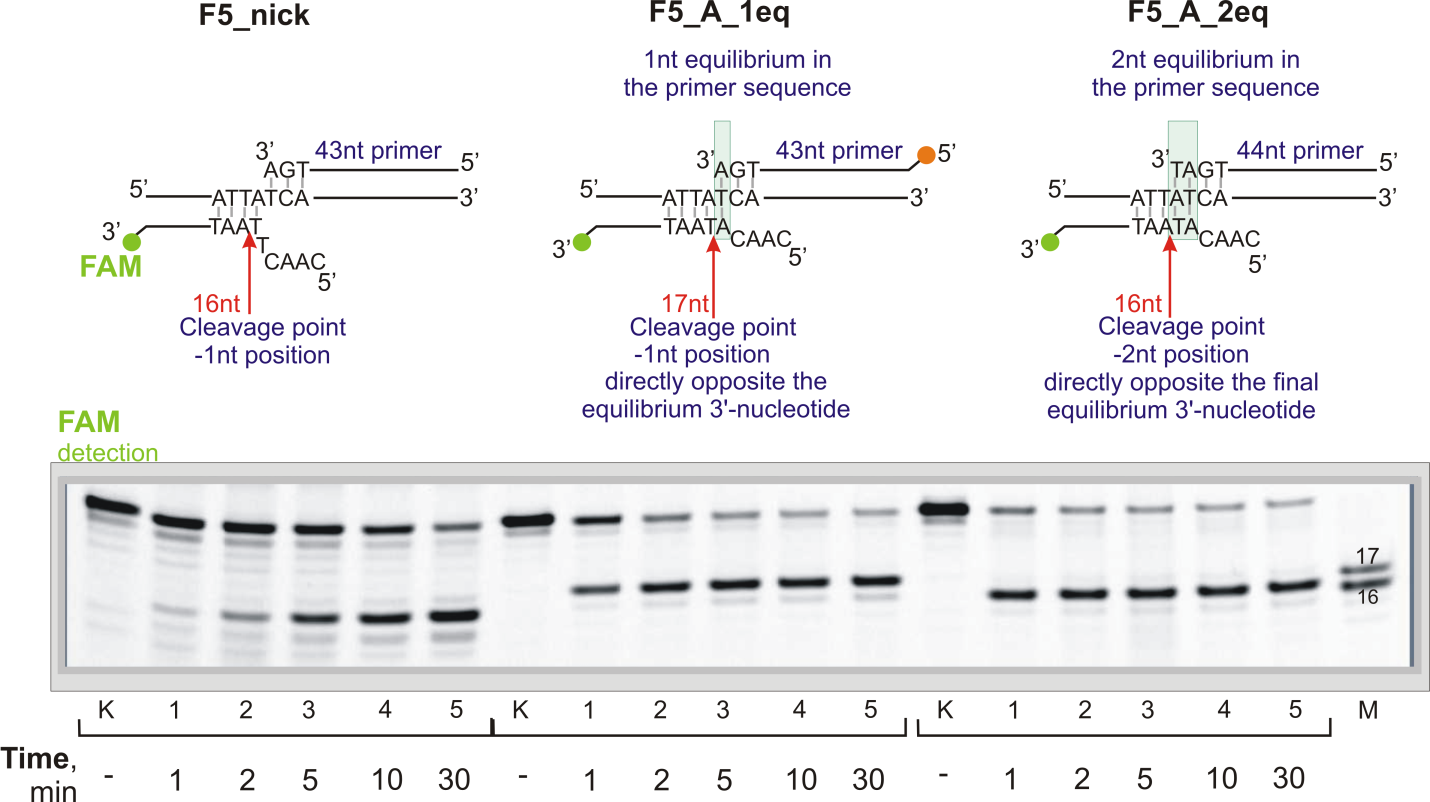

**Figure S8.** The cleavage point is located directly opposite the terminal equilibrium 3′-nucleotide of the primer. Without equilibrium nucleotides (F5-nick) and with a 1nt equilibrium area (F5_A_1eq), FEN1 cleaves at the (-1nt) position. With a 2nt equilibrium area (F5_A_2eq), FEN1 cleaves at the (-2nt) position.

Schematic representations of the substrates are depicted at the top; the lime green circle indicates the FAM-label position; the green arrow indicates the position of the substrate DNA. Lane “k” is the DNA control; lane “M” is the length marker. Reaction mixtures contained 20 nM DNA and 20 nM FEN1.

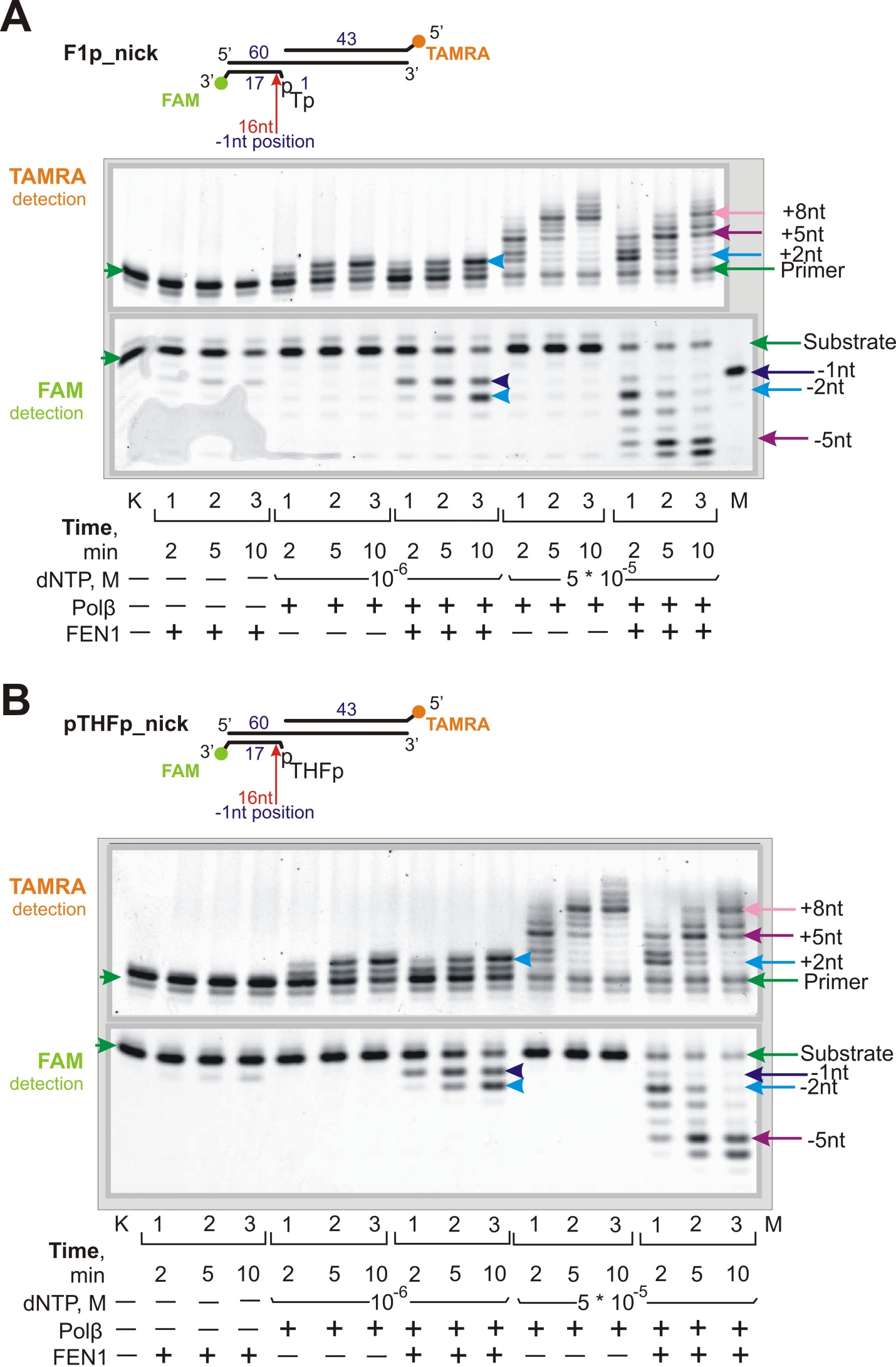

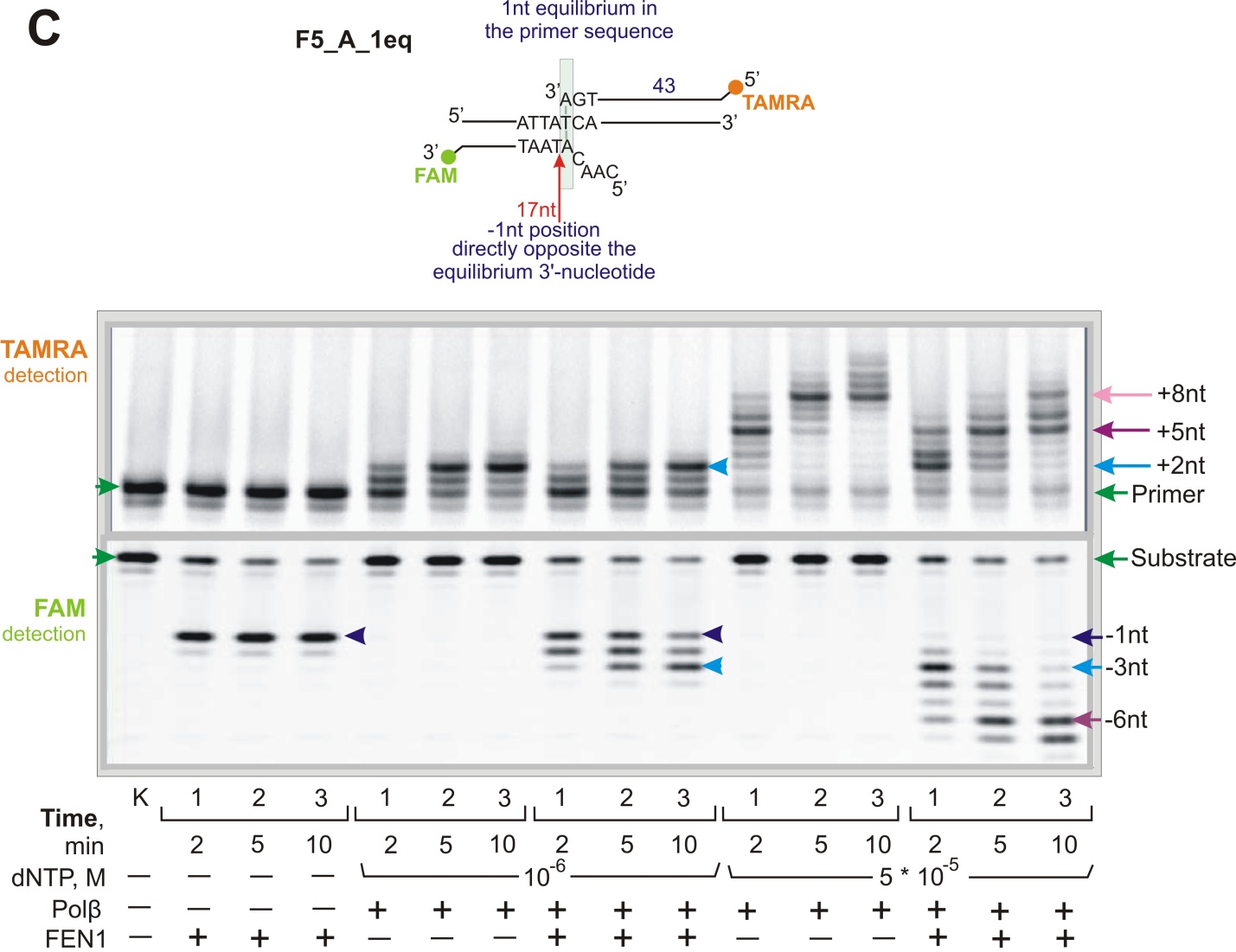

**Figure S9.** The cleavage product profiles for F1p_nick (**A**), pTHFp-nick (**B**), and F5_A_1eq (**C**) DNA correspond to the primer extension products at 1 and 50 µM dNTP concentrations.

The names and schematic representations of the substrates are depicted above the gels; the lime green and orange circles indicate FAM and TAMRA labels, respectively; the green arrows indicate the position of the substrate DNA (primer or flap-strand); the light blue, dark blue, and plum arrows indicate primer elongation and the corresponding cleavage products; lane “k” is the DNA control, and “M” is the length marker. Reaction mixtures contained 20 nM DNA, 20 nM FEN1, 20 nM Polβ, and dNTPs at the indicated concentration.

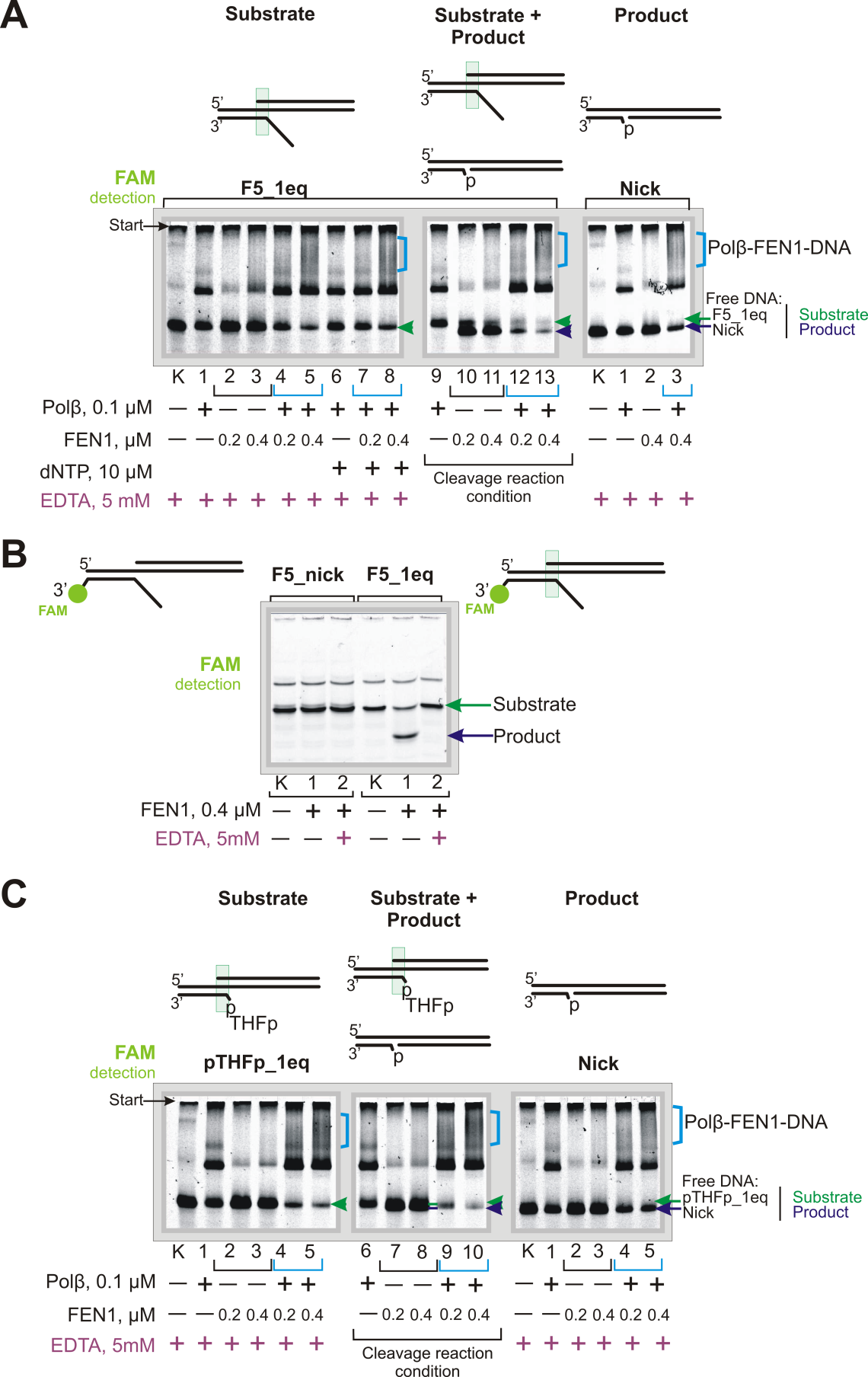

**Figure S10.** The Polβ-FEN1-DNA ternary complex was detected using FEN1 substrate DNAs (pTHFp_1eq and F5_1eq) and product Nick DNA.

**A**.	The ternary complex is registered with the 5nt-flap DNA containing one equilibrium nucleotide, in the presence of 5mM EDTA (lanes 1-8) and in the absence of EDTA (lanes 9-13). Lanes 9-13 correspond to the cleavage reaction conditions.

**B.**	FEN1 is able to cleave F5_1eq DNA without the addition of magnesium/manganese (lane 1) since these metal ions can remain bound within the protein structure while obtaining during the purification process. The addition of 5 mM EDTA leads to the disappearance of cleavage products (lane 2 for F5_1eq). The F5_nick DNA structure, without equilibrium nucleotides, was not cleaved without the addition of magnesium/manganese. The reaction products were separated by denaturing gel electrophoresis (20% UREA-PAGE (acrylamide/*bis*-acrylamide = 19:1, 8 M urea, and 10% or 20% formamide)).

**C.**	The ternary complex is also formed with pTHFp_1eq DNA containing one equilibrium nucleotide, in the presence of 5mM EDTA (lanes 1-5) and in its absence (lanes 6-10). Lanes 6-10 correspond to the cleavage reaction conditions.

A schematic representation of the substrates is depicted at the top; the green arrow indicates the position of free, unbound substrate DNA, and the dark blue arrow indicates the free product DNA; the blue staples indicate putative Polβ-FEN1-DNA complexes. Lane “k” is the DNA control. Reaction mixtures (10 μL) contained 20 nM DNA, 100 nM Polβ, 5 mM EDTA, 10 µM dNTPs, and FEN1 at concentrations of 200 or 400 nM.

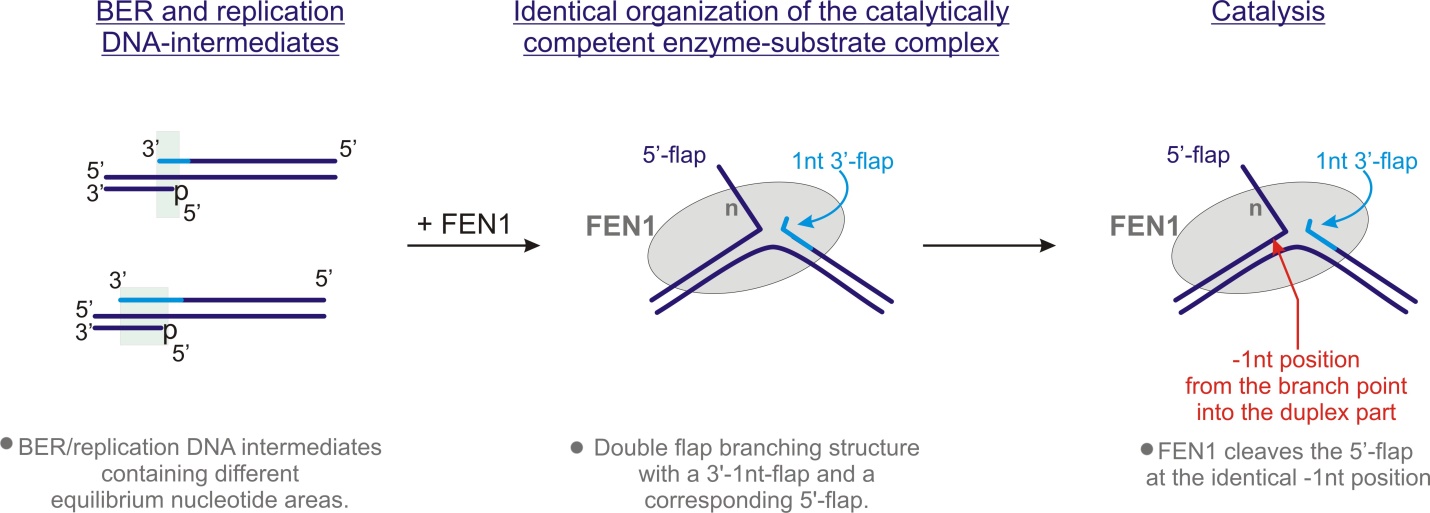

**Figure S11.** FEN1’s structural specificity is provided by the identical organization of the enzyme-substrate complex and is independent of the length of the equilibrium nucleotide area in the DNA substrate. FEN1 binds to the 3′-primer nucleotide and forms a double-flap branching structure with a 1nt 3′- flap and a corresponding 5′-flap. FEN1 then cleaves the 5′-flap in the bound DNA-intermediate at the (-1nt) position from the branch point into the duplex part.
